# A Trem2*R47H mouse model without cryptic splicing drives age- and disease-dependent tissue damage and synaptic loss in response to plaques

**DOI:** 10.1101/2022.03.09.483490

**Authors:** Kristine M. Tran, Shimako Kawauchi, Enikö A. Kramár, Narges Rezaie, Heidi Yahan Liang, Miguel Arreola, Celia Da Cunha, Jimmy Phan, Sherilyn Collins, Amber Walker, Jonathan Neumann, Giedre Milinkeviciute, Angela Gomez-Arboledas, Dominic I. Javonillo, Katelynn Tran, Magdalena Gantuz, Stefania Forner, Vivek Swarup, Andrea J. Tenner, Frank LaFerla, Marcelo A. Wood, Ali Mortazavi, Grant R. MacGregor, Kim N. Green

## Abstract

Genome-Wide Association Studies revealed the *TREM2* R47H variant as one of the strongest genetic risk factors for late-onset Alzheimer’s Disease (AD). Unfortunately, many current *TREM2*\*R47H mouse models are associated with cryptic mRNA splicing of the mutant allele that produces a confounding reduction in protein product. We have developed the *Trem2*^R47H NSS^ (<u>N</u>ormal <u>S</u>plice <u>S</u>ite) mouse model where the *Trem2* allele is expressed at a similar level to the wild-type *Trem2* allele, without evidence of cryptic splicing products, and appropriate inflammatory responses to cuprizone challenge. Utilizing the 5xFAD mouse model, we report age- and disease-dependent changes in response to pathology. At an early disease stage (4 mo), homozygous *Trem2*^R47H NSS^; hemizygous 5xFAD (*Trem2*^R47H NSS^ ; 5xFAD) mice have reduced size and number of microglia plus impaired interaction with plaques, that is associated with increased dystrophic neurites and axonal damage detected through plasma neurofilament light chain (NfL) level and suppressed inflammation. However, homozygosity for *Trem2*^R47H NSS^ suppressed LTP deficits and presynaptic puncta loss caused by the 5xFAD transgene array. At a more advanced disease stage (12 mo,) *Trem2*^R47H NSS^ ; 5xFAD mice no longer display impaired plaque-microglia interaction or suppressed inflammatory gene expression, although NfL levels remain elevated, and a unique interferon-related gene expression signature is seen. Furthermore, *Trem2*^R47H NSS^ ; 5xFAD mice also display robust LTP deficits and exacerbated presynaptic loss. Collectively, we provide a *Trem2*^R47H^ variant mouse without cryptic splicing, and demonstrate it has disease stage dependent effects when combined with a plaque bearing model, with an initial loss of function that ultimately resolves, giving rise to a unique interferon signature and associated tissue damage.

## INTRODUCTION

Triggering receptor expressed on myeloid cells 2 (TREM2) is a cell surface receptor expressed in myeloid cells that participates in sensing of the environment surrounding microglia. Mutations / variants within *TREM2* are associated with age-dependent development of several neurodegenerative diseases including Alzheimer’s disease (AD), Frontotemporal dementia (FTD), Parkinson’s disease (PD), Amyotrophic lateral sclerosis (ALS), and Nasu-Hakola disease, depending on the specific variant. Loss of function TREM2 mutations result in Nasu-Hakola disease, a leukodystrophy with similarities to adult-onset leukoencephalopathy with axonal spheroids and pigmented glia (ALSP) that is caused by loss of function mutations in colony-stimulating factor-1 receptor (CSF1R). TREM2 and CSF1R both converge on the same intracellular pathways in myeloid cells mediated by DNAX-activating protein of 12kDa (DAP12, also known as TYROBP), in which mutations also lead to Nasu-Hakola disease (Otero et al., 2009; Paloneva et al., 2002). Other *TREM2* mutations likely modify protein function and increase risk for the development of these other neurodegenerative diseases, resulting in varied pathologies in distinct brain regions. Thus, understanding TREM2 biology and the roles that it plays in brain homeostasis is critical, as its dysfunction or actions can manifest as leukodystrophies or predominantly grey matter targeting pathologies.

The R47H missense variant in *TREM2* is strongly and reproducibly linked to the development of Late-Onset AD (LOAD; (Guerreiro et al., 2013; Jonsson et al., 2013)). TREM2 is a membrane spanning protein with an extracellular ectodomain (aa’s 1-174), and a shorter intracellular sequence (aa’s 196-230), which interacts with DAP12 and regulates gene expression and other pathways via immunoreceptor tyrosine-based activation motifs (ITAM; (Linnartz et al., 2010)). The R47 residue is located within the extracellular Ig-like domain portion, and likely modifies interactions of TREM2 with its ligands (Song et al., 2017; Wang et al., 2015), which include phospholipids, HDL, LDL, APOE, APOJ (clusterin), sulfatides, bacterial lipopolysaccharide, DNA, apoptotic neurons and Ab. Notably, while the R47H variant is associated with AD, it is also associated with increased risk of development of FTD, PD, and ALS (Borroni et al., 2014; Cady et al., 2014; Rayaprolu et al., 2013). Given that the same alteration in TREM2 function can manifest as distinct disease with distinct pathologies (affecting discrete brain regions) and clinical domains, we hypothesize that altered TREM2 function provides more permissive conditions for disease progression (e.g., an inadequate immune response to pathogenic species), rather than being a specific trigger for disease onset. For example, in LOAD, plaques develop many years prior to clinical symptoms and neurodegeneration, which correlate with the spread of Tau not amyloid pathology. Thus, it is plausible that mutations/variants in *TREM2* may modulate the brains’ response (or lack of response) to plaques, ultimately driving neurodegeneration. Similar reactions to pathologies may occur in PD, FTD, and ALS, although the association with these other diseases have failed to replicate in some studies (Lill et al., 2015; Zhang et al., 2020).

Microglia are the primary immune cells of the central nervous system and play important roles in responding to pathological insults and maintaining tissue homeostasis. Given that *TREM2* is primarily expressed by microglia, these cells have been strongly implicated in the development of LOAD, as well as other TREM2-associated neurological disorders. Several non-coding polymorphisms in myeloid expressed genes have been identified via GWAS and linked to LOAD. These include *SPI1, BIN1, GRN, CD33* (Carrasquillo et al., 2010; Jiang et al., 2014; Lambert et al., 2013; Tansey et al., 2018)), as well as coding variants in *ABI3* and *PLCG2* (Conway et al., 2018; Sims et al., 2017). During AD, microglia mount an inflammatory response to Aβ plaques, as evidenced by findings in both human AD brains and animal models of the disease (Leng and Edison, 2021; Sayed et al., 2021). Accumulating evidence implicates microglia in several AD-related processes including plaque formation and growth (Spangenberg et al., 2019), plaque compaction (Casali et al., 2020; Spangenberg et al., 2019), constituting a protective barrier against dystrophic neurites (Condello et al., 2015), promoting or preventing development and spreading of Tau pathology (Shi et al., 2019), cerebral amyloid angiopathy (Spangenberg et al., 2019), destruction of perineuronal nets (Arreola et al., 2021; Crapser et al., 2020), as well as synaptic and neuronal loss (Arreola et al., 2021; Elmore et al., 2018; Hong et al., 2016; Rice et al., 2015; Schafer et al., 2012; Spangenberg et al., 2016; Wang et al., 2020; Werneburg et al., 2020).

Despite these findings, it remains unclear whether the microglial response to plaques protects against or promotes disease progression. *Trem2* KO mice have microglia that fail to respond appropriately to plaques resulting in a lack of an inflammatory cascade or physical association with plaques, and this appears to accelerate aspects of disease progression (Wang et al., 2016). On the other hand, *Tyrobp* KO mice also have microglia that fail to respond to plaques, but this appears to be protective (Haure-Mirande et al., 2017). As TREM2 loss of function in human results in Nasu-Hakola disease, its absence in mice may have unintended consequences when explored in the context of plaques. In support of a beneficial inflammatory-mediated response, the PLCG2*P522R coding variant enhances microglial-evoked inflammation in cultures and is associated with reduced risk of developing LOAD (Kleineidam et al., 2020) while suppression of inflammation and increased risk of developing LOAD is associated with PLCG2*M28L variant (Tsai et al.,2020). Microglial depletion in mouse models of AD assert that disease stage and timing are important for microglial function in AD, with protective roles early on, but damaging roles at later stages of disease (Spangenberg et al., 2019; Spangenberg et al., 2016). Establishing which aspects of microglial biology promote vs retard disease progression is critical for clinical success with microglial targeting strategies. To date, conflicting *in vivo* data, utilizing Trem2*R47H mouse models have made it difficult to resolve TREM2 function in the context of both amyloid and Tau pathologies. For example, several studies with *Trem2*\*R47H models indicated that it acts as a loss of function and demonstrates impaired association and reaction to plaques (Cheng-Hathaway et al., 2018; Xiang et al., 2018), and yet associates with an increased risk of disease. However, subsequent findings indicated that generation of *Trem2*\*R47H mouse models via CRISPR resulted in an unintended induction of cryptic splicing products in the mutant *Trem2* allele, resulting in dramatically reduced TREM2 protein, which is not observed in human R47H carriers (Xiang et al., 2018).

Here, we developed a *Trem2*^R47H^ mouse model that does not display cryptic splicing and has normal level of transcription of the *Trem2*^R47H^ allele. We show that the TREM2*R47H variant does not act as a loss of function allele. We demonstrate that when combined with the 5xFAD mouse model of AD (Oakley et al., 2006), homozygosity for the TREM2*R47H variant causes an initial suppression of microglial plaque response, which at later stages is lost, and even exacerbated, inducing an interferon-related signature – indicating an age-dependent effect of TREM2 dysfunction on microglial biology and AD pathology. Notably, homozygosity for the TREM2*R47H variant promotes tissue damage in response to plaques, as assessed by increased dystrophic neurites, and brain and plasma neurofilament light chain (NfL) levels. Furthermore, homozygosity for the TREM2*R47H variant results in marked synaptic loss and LTP deficits by 12 months of age, highlighting the detrimental impact of dysfunctional microglia and TREM2 on neurons.

## METHODS

### Animals

All experiments involving mice were approved by the UC Irvine Institutional Animal Care and Use Committee and were conducted in compliance with all relevant ethical regulations for animal testing and research. All experiments involving mice comply with the Animal Research: Reporting of *In Vivo* Experiments (ARRIVE) guidelines, which are specifically addressed in the Supplementary Materials.

### Mice

To generate *Trem2*^R47H^ mice (B6(SJL)-*Trem2^em1Aduci^*/J, Stock number #034036 -JAX), Alt-R Crispr RNA’s (TMF1342 – gaagcactgggggagacgca, TMF-1343 – gtacatgacaccctcaagga) and tracrRNA plus CAS9 protein (IDT) as a ribonucleoprotein (RNP) was microinjected into C57BL/6J zygotes along with a ssODN sequence (TMF1341 – sequence available upon request) to introduce the R47H missense mutation. G0 founder animals containing the desired DNA sequence changes were backcrossed with C57BL/6J mice and N1 heterozygous mice were sequenced to determine the mutant allele. N1 heterozygous mice were backcrossed to produce N2F1 heterozygotes, which were used for subsequent analysis. *Trem2*^R47H^ heterozygous mice were then bred to generate homozygous *Trem2*^R47H^ animals used in the study. 5xFAD hemizygous (B6.CgTg(APPSwFlLon,PSEN1*M146L*L286V)6799Vas/Mmjax, Stock number 34848-JAX, MMRRC) and non-transgenic littermates were produced by natural mating or IVF procedures with C57BL/6J (Jackson Laboratory, ME) females. After weaning, they were housed together with littermates and aged until the harvest dates. All animals were bred by the Transgenic Mouse Facility at UCI.

### Genotyping

For *Trem2*^R47H^ genotyping, we used a common primer set to amplify both *Trem2* wildtype allele and *Trem2*^R47H^ allele (For 5′-TCAACACCACGGTGCT -3′ and Rev 5′-TGTGTGCTCACCACACG

-3′). Two fluorophore labeled-hydrolysis probes which hybridized specific to mouse *Trem2* wildtype amplicon (5’-TGCGTCTCCCCCAGTGCTTCAA-3’+HEX) and *Trem2* R47H mutation (5’- TATGTCTGCCCCAATGTTTCAACGCG-3’-FAM) were used to detect the allelic ratio in the amplicon. The relative fluorescence units from each probe at the end point of PCR cycles were plotted to call the genotype by using Allelic Discrimination function using Bio-Rad CFX Maestro software. For 5xFAD genotyping, hydrolysis probe which hybridizes APP(Swe) mutation amplicon was used (For 5′-TGGGTTCAAACAAAGGTGCAA-3′ and Rev 5′- GATGACGATCACTGTCGCTATGAC-3′: APP(Swe) probe 5′-CATTGGACTCATGGTGGGCGGTG-3′.) to detect transgenes. We used the endogenous *ApoB* allele (For 5′-CACGTGGGCTCCAGCATT-3′ and Rev 5′-TCACCAGTCATTTCTGCCTTTG-3′: *ApoB* probe 5′-CCAATGGTCGGGCACTGCTCAA-3′) to normalize the Ct values.

### Cuprizone (CPZ) animal treatment

Eight-week-old C57BL/6J, *Trem2*^em1Aduci^, *Trem2*^em1Adiuj^, and *Trem2*^em2Adiuj^ mice (Stock number: 000664, 034036, 027918, 027197 respectively; The Jackson Laboratory) were used. Each mouse model had 2 groups of 4 male mice: Control on 6 weeks standard chow, and CPZ on 6 weeks 0.3% cuprizone chow (Envigo). Mice within the same experiment group (i.e. same genotype and diet) were housed together for the duration of feeding. Weights of individual mouse and chow consumptions of each cage were recorded, and chow were changed every 3 or 4 days to monitor expected weight loss as well as ensuring freshness of cuprizone chow. Brains were collected and fixed in 4% paraformaldehyde for 24 hr followed by cryoprotection by immersion in 5% sucrose for 24 h then 30% sucrose for 5 days.

### Histology

Mice were euthanized at 4 and 12 month of age via CO2 inhalation and transcardially perfused with 1X phosphate buffered saline (PBS). For all studies, brains were removed and hemispheres separated along the midline. Brain halves were either flash frozen for subsequent biochemical analysis or drop-fixed in 4% paraformaldehyde (PFA (Thermo Fisher Scientific, Waltham, MA)) for immunohistochemical analysis. Fixed half brains were sliced at 40 μm using a Leica SM2000R freezing microtome. All brain hemispheres were processed and coronal brain slices (between - 2.78mm posterior and –3.38mm posterior to Bregma according to the Allen Mouse Brain Atlas, Reference Atlas version 1, 2008) imaged via a Zeiss Axio Scan Z1 Slidescanner using a 10X 0.45 NA plan-apo objective. Images were also acquired using a 20X objective via a Leica TCS SPE-II confocal microscope and quantified using Bitplane Imaris Software.

### Immunohistochemistry

One representative brain slice from each mouse of the same experimental group (i.e. same genotype, age, and sex) was stained in the same well. Free-floating sections were washed three times with 1X PBS (1 x 10 min and 2 x 5 min) and for Thiosflavin-S staining, 10 min incubation in 0.5% Thio-S (1892; Sigma-Aldrich, St. Louis, MO) diluted in 50% ethanol followed. Sections were then washed 2X for 5 min each in 50% ethanol and one 10-min wash in 1xPBS. For Amylo-Glo staining, following the previously described PBS washes, the free-floating brain slices were washed in 70% ethanol for 5 min and rinsed in deionized water for 2 min before being immersed for 10 min in Amylo-Glo RTD Amyloid Plaque Staining Reagent (1:100; TR-200- AG; Biosensis, Thebarton, South Australia) diluted in 0.9% saline solution. Afterwards, sections were washed in 0.9% saline solution for 5 min then rinsed in deionized water for 15 sec before proceeding with a standard indirect immunohistochemical protocol. From the incubation period for both Thio-S and Amylo-Glo onwards, sections were kept under foil or in the dark. Sections were immersed in normal blocking serum solution (5% normal goat serum with 0.2% Triton X-100 in 1X PBS) for 1 hr before overnight incubation at 4°C in primary antibodies diluted in normal blocking serum solution.

Brain sections were stained following a standard indirect technique as described (Forner et al., 2021; Javonillo et al., 2021) with the following primary antibodies against: ionized calcium-binding adapter molecule 1 (IBA1; 1:2000; 019-19741; Wako, Osaka, Japan), Aβ1-16(6E10; 1:2000; 8030001; BioLegend, San Diego, CA), glial fibrillary acidic protein (GFAP; 1:1000; AB134436; Abcam, Cambridge, MA), S100β (1:200; AB41548; Abcam, Cambridge, MA), lysosome-associated membrane protein 1 (LAMP1; 1:200; AB25245; Abcam, Cambridge, CA), neurofilament light chain (NfL; 1:200; 171 002; Synaptic Systems, Germany), CD74 (1:500; 151002 ; BioLegend, San Diego, CA), CD11c (1:100.; 50-112-2633; eBioscience), myelin basic protein (MBP; 1:200; MAB386; Milipore Sigma), Synaptophysin (1:1000; Sigma-Aldrich), PSD-95 (1:500; Abcam and Cell Signaling).

High-resolution fluorescence images were obtained using a Leica TCS SPE-II confocal microscope and LAS-X software. For confocal imaging, one field of view (FOV) per brain region was captured per mouse using the Allen Brain Atlas to capture comparable brain regions.

### Soluble and insoluble fraction A**β** levels

Preparation of samples and quantification of Ab was performed as described (Forner et al., 2021; Javonillo et al., 2021). Micro-dissected hippocampal and cortical regions of each mouse were flash-frozen and processed for biochemical analysis. Samples were pulverized using a Bessman Tissue Pulverizer kit. Pulverized hippocampal tissue separated for biochemical analysis was homogenized in 150µL of Tissue Protein Extraction Reagent (TPER; Life Technologies, Grand Island, NY), while cortical tissue was homogenized in 1000µL/150 mg of TPER. This composition of TPER includes 25mM bicine and 150mM sodium chloride (pH 7.6) to efficiently solubilize proteins within brain tissue following homogenization. Together with protease (Roche, Indianapolis, IN) and phosphatase inhibitors (Sigma-Aldrich, St. Louis, MO), the homogenized samples were centrifuged at 100,000 g for 1 hr at 4°C to generate TPER-soluble fractions. For formic acid-fractions, pellets from TPER-soluble fractions were homogenized in 70% formic acid: 75µL for hippocampal tissue or half of used TPER volume for cortical tissue. Afterwards, samples were centrifuged again at 100,000 g for 1 hr at 4°C. Protein in the insoluble fraction of micro- dissected hippocampal and cortical tissue were normalized to its respective brain region weight, while protein in soluble fractions were normalized to the protein concentration determined via Bradford Protein Assay. Formic acid neutralization buffer was used to adjust pH prior to running ELISAs.

Quantitative biochemical analyses of human Aβ soluble and insoluble fraction levels were acquired using the V-PLEX Aβ Peptide Panel 1 (6E10) (K15200G-1; Meso Scale Discovery, Rockville, MD). Finally, quantitative biochemical analysis of neurofilament-light chain (NfL) in plasma was performed using the R-Plex Human Neurofilament L Assay (K1517XR-2; Meso Scale Discovery, Rockville, MD).

### Imaris quantitative analysis

Confocal images of each brain region were quantified automatically using the spots module within the Imaris v9.7 software (Biplane Inc. Zürich, Switzerland) then normalized to the area of the field-of-view (FOV). Amyloid burden was assessed by measuring both the total Thio-S^+^ plaque number normalized to FOV area and their volume via the surfaces module in Imaris software. Similarly, volumetric measurements (i.e. Thio-S^+^ plaque volume, IBA1^+^ microglia volume, etc.) were acquired automatically utilizing the surfaces module on confocal images of each brain region. Quantitative comparisons between experimental groups were carried out in sections stained simultaneously.

For synaptic quantifications, three FOVs at 63x objective per brain region were captured, and quantifications for each animal were averaged. Total cell counts and morphological analyses were obtained by imaging comparable sections of tissue from each animal using a 20× objective, at multiple z-planes, followed by automated analyses using Bitplane Imaris 7.5 spots and filaments, respectively, as described (Forner et al., 2021; Javonillo et al., 2021). Co-localization analyses were conducted using Bitplane Imaris 7.5 colocalization and surfaces modules. For hemisphere stitches, automated slide scanning was performed using a Zeiss AxioScan.Z1 equipped with a Colibri camera and Zen AxioScan 2.3 software. Cell quantities were determined using the spots module in Imaris.

### Long-term potentiation

Hippocampal slices were prepared from WT, *Trem2*^R47H^, 5xFAD, 5xFAD/ *Trem2*^R47H^ (8-10 mice per sex per genotype) at 4 and 12 months of age. Hippocampal slice preparation and long-term potentiation (LTP) recording was performed as described (Forner et al., 2021; Javonillo et al., 2021). Following isoflurane anesthesia, mice were decapitated and the brain was quickly removed and submerged in ice-cold, oxygenated dissection medium containing (in mM): 124 NaCl, 3 KCl, 1.25 KH2PO4, 5 MgSO4, 0 CaCl2, 26 NaHCO3, and 10 glucose. Coronal hippocampal slices (340µm) were prepared using a Leica vibrating tissue slicer (Model: VT1000S) before being transferred to an Interface recording containing preheated artificial cerebrospinal fluid (aCSF) of the following composition (in mM): 124 NaCl, 3 KCl, 1.25 KH2PO4, 1.5 MgSO4, 2.5 CaCl2, 26 NaHCO3, and 10 glucose and maintained at 31±1°C. Slices were continuously perfused with this solution at a rate of 1.75–2ml/min while the surface of the slices were exposed to warm, humidified 95% O2 / 5% CO2. Recordings began following at least 2 hr of incubation. Field excitatory postsynaptic potentials (fEPSPs) were recorded from CA1b striatum radiatum using a single glass pipette filled with 2M NaCl (2–3 MΩ) in response to orthodromic stimulation (twisted nichrome wire, 65 µm diameter) of Schafer collateral-commissural projections in CA1 striatum radiatum. Pulses were administered at 0.05Hz using a current that elicited a 50% maximal response. Paired-pulse facilitation was measured at 40, 100, and 200sec intervals prior to setting baseline. After establishing a 20 minute stable baseline, the orthodromic stimulated pathway was used to induce LTP by delivering 5 ‘theta’ bursts, with each burst consisting of four pulses at 100Hz and the bursts themselves separated by 200 msec (i.e., theta burst stimulation or TBS). The stimulation intensity was not increased during TBS. Data were collected and digitized by NAC 2.0 Neurodata Acquisition System (Theta Burst Corp., Irvine, CA) and stored on a disk.

### RNA sequencing

RNA sequencing was performed as described (Forner et. al., 2021). Total RNAs were extracted using RNeasy Mini Kit (Qiagen) on a QIAcube (Qiagen) liquid handling platform. RNA integrity number (RIN) was measured by Qubit RNA IQ Assay (Invitrogen) and samples with RIN >= 7.0 were kept for cDNA synthesis. cDNA synthesis and amplification were performed followed by Smart-seq2 [Picelli, S., et al. Nature Protocols. 2014. 9(1): p. 171.] standard protocol.

Libraries were constructed by using the DNA Prep Kit (Illumina) on an epMotion 5070 TMX (Eppendorf) automated pipetting system. Libraries were base-pair selected based on Agilent 2100 Bioanalyzer profiles and normalized determined by KAPA Library Quantification Kit (Roche). The libraries were built from 3 to 5 different mice per genotype, sex and tissue (hippocampus) across 2 different timepoints (4, and 12 months). For cuprizone experiment, whole brains were used for 4 males per genotype per condition (control or CPZ) at 12 months of age. The 4 month of mouse libraries were sequenced using paired-end 43bp mode on Illumina NextSeq500 platform with >14 million reads per sample. The 12 month old mouse libraries were sequenced using paired-end 101bp mode on Illumina NextSeq2000 platform with > 28 million reads per sample. Sequences were aligned to the mouse genome (mm10) and annotation was done using GENCODE v21. Reads were mapped with STAR v.2.7.3a and RSEM (v.1.3.3) was used for quantification of gene expression.

#### Differential gene expression analysis

Differential gene expression analysis was performed using edgeR per timepoint and genotype. Genes with a log2(Fold Change) >1 and various threshold for value depending comparison were labelled. To compare different sets of genes differentially expressed we created a binary matrix identifying up and downregulated genes across different comparisons. A matrix indicating up or downregulation was later used to plot a heatmap. From the comparisons, lists of genes of interest were chosen to plot a heatmap of their expression and a GO term enrichment analysis using enrichR (https://amp.pharm.mssm.edu/Enrichr/) and the top 4 GO terms were plotted.

#### Weighted correlation gene network analysis analysis

Weighted gene correlation network analysis (WGCNA) was used on two different datasets: 1.*TREM2* dataset with matching wildtype (5xFAD) in 2 different timepoint (4 month and 12 month) in hippocampus in both sex 2. Cuprizone and control group in male brain samples at 12 month. For both datasets we used a matrix filtered by genes with more than 1 TPM and without an outlier sample. Based on our datasets we used power 15 as a soft threshold for first dataset and power 5 for second one. The other parameters were same for both including: min. module size =50 and MEDissThres = 0.2.We identified significant modules by calculating the correlation with the traits, then proceeded to plot the behavior per sample of the genes in the red, royal blue module (first dataset) and turquoise, cyan and royalblue module (second dataset), by doing a GO term analysis using Metascape (https://metascape.org).

### Statistics

Every reported *n* represents the number of independent biological replicates. The sample sizes are similar with those found in prior studies conducted by MODEL-AD and were not predetermined using statistical methods (Forner et al., 2021; Javonillo et al., 2021). Electrophysiology, immunohistochemical, and biochemical data were analyzed using Student’s t-test, one-way ANOVA, or two-way ANOVA via Prism v.9 (GraphPad, La Jolla, CA). Bonferroni-Šídák and Tukey’s post hoc tests were utilized to examine biologically relevant interactions from the two-way ANOVA. Where sex-differences are apparent, a Student’s t-test was used within genotype group. *p ≤ 0.05, ** p ≤ 0.01, ***p ≤ 0.001, ****p ≤ 0.0001. Statistical trends are accepted at p < 0.10 (^#^). Data are presented as raw means and standard error of the mean (SEM).

## RESULTS

### *The Trem2*^R47H NSS^ mutation promotes loss of oligodendrocyte gene expression in response to cuprizone treatment

Coding variants strongly linked to Late-Onset AD (LOAD) offer an invaluable resource to understand the biology that leads to disease pathogenesis and progression. The R47H missense mutation in *TREM2* directly implicates microglial biology in LOAD, and elucidating the underlying mechanisms depends on experimental models that recapitulate the human function. Results of previous studies of mice with the *TREM2* R47H missense mutation introduced via CRISPR suggested that it acts as a near-complete loss of function, recapitulating phenotypes seen in *Trem2* KO mice (Cheng-Hathaway et al., 2018; Kotredes et al., 2021). However, subsequent analyses of *Trem2* expression and splicing in these models identified the unexpected generation of a cryptic splice site and subsequent loss of *Trem2* expression due to the synonymous codon changes introduced as part of the CRISPR repair template (Xiang et al., 2018). Given the importance of a *Trem2*\*R47H mouse model that more accurately recapitulates human cases, we designed alternative CRISPR repair templates, guided in part by (Cheng et al., 2018), to introduce the R47H mutation into C57BL/6J mice. Confirmation of sequence change (CGC – CAT; arginine – histidine) and synonymous silent codon changes are shown in Fig. 1a and mapping of reads from bulk-RNA-seq from the brains of wild-type and homozygous to the *Trem2* locus showed no evidence of unusual splicing events. We designated the model as *Trem2*^R47H NSS^ (<u>Normal Splice Site</u>; available at The Jackson Laboratory - stock #034036). To assess the impact of *Trem2*^R47HNSS^ on inflammation and to test if it acts as a loss of function allele, we utilized a cuprizone model of demyelination. We included cohorts of *Trem2*^R47H^ mice with the identified <u>Cryptic Splice Site</u> and reduced expression ((Xiang et al., 2018); designated *Trem2*^R47H CSS^), and *Trem2* KO mice. These 3 groups, along with a wild-type group were treated with cuprizone (0.3%) or control in chow for 6 weeks (Fig. 1b), then were sacrificed and brains examined by histology and RNA-seq. Whole brain *Trem2* expression values were plotted, showing that *Trem2*^R47H NSS^ mice have similar *Trem2* expression levels to wild-type mice, and that both *Trem2*^R47H CSS^ and *Trem2* KO mice have reduced expression (Fig. 1c). Notably, with cuprizone, *Trem2* levels increased similarly in wild-type and *Trem2*^R47H NSS^ mice, and to a lesser extent in *Trem2*^R47H CSS^ mice. No aberrant splicing was observed in *Trem2*^R47H NSS^ mice (data not shown).

**Figure 1:**
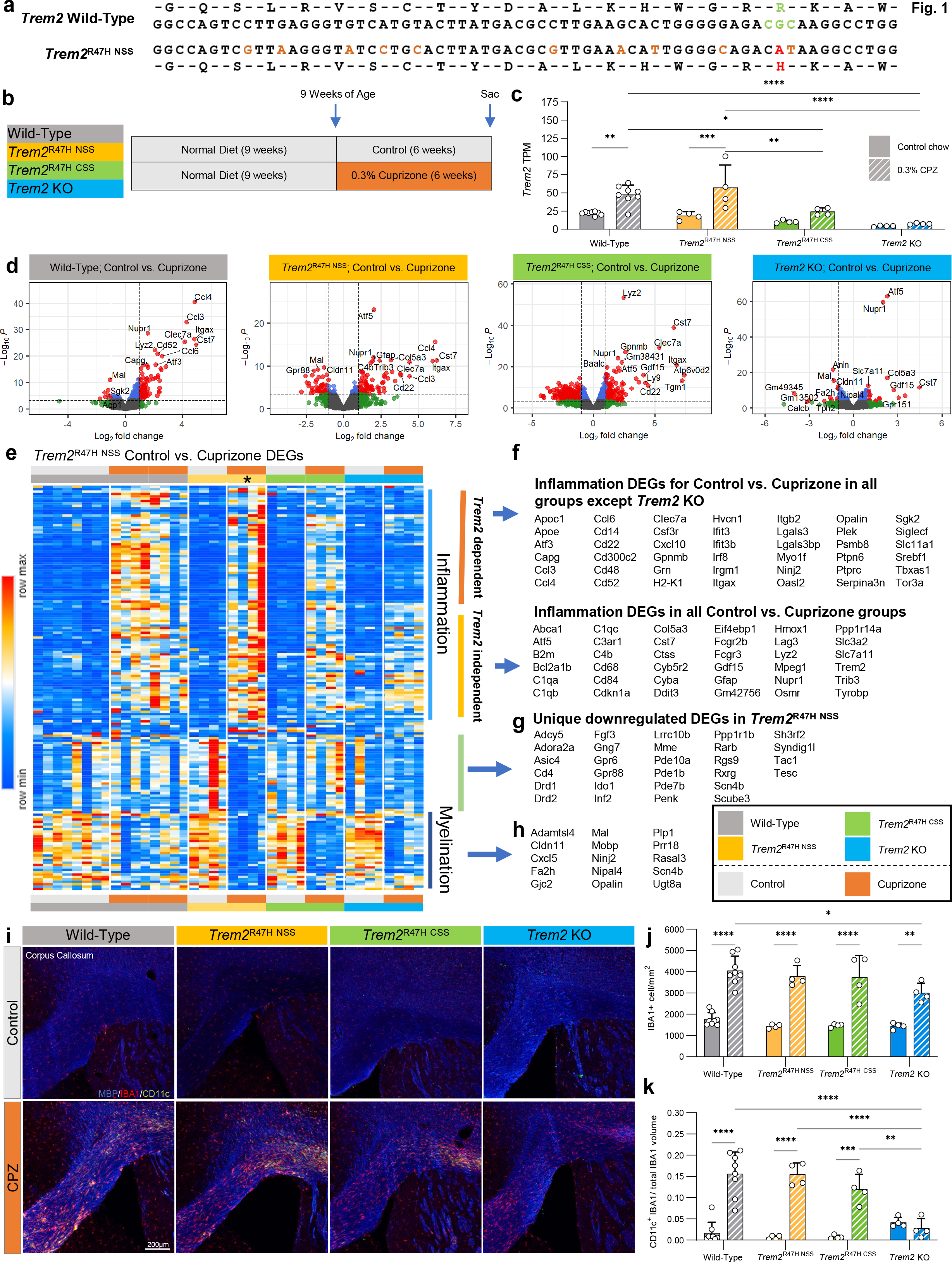
Cuprizone model of demyelination on wild-type, *Trem2*^R47H NSS^, *Trem2*^R47H CSS^, and *Trem2* KO mice. **a** The codon sequence for arginine is shown in green with the G to A transition that encodes histidine (H) shown in red in the *Trem2*^R47H NSS^ allele. Ten silent DNA mutation are shown in tan. **b** Cuprizone feeding scheme of wild-type, *Trem2*^R47H NSS^, *Trem2*^R47H CSS^ and *Trem2* KO. **c** *Trem2* TPM values of wild-type, *Trem2*^R47H NSS^, *Trem2*^R47H CSS^ and *Trem2* KO males examined from bulk, whole-brain RNA sequencing data showed increase in *Trem2* expression in response to cuprizone treatment in wild-type and *Trem2*^R47H NSS^ but not in *Trem2*^R47H CSS^ or *Trem2* KO mice, highlighting similar *Trem2* expression level between wild-type and *Trem2*^R47H NSS^ but not *Trem2*^R47H CSS^ and *Trem2* KO. **d** Volcano plot of differentially expressed genes, displaying fold change of genes (log2 scale) and *P* values (−log10 scale) between control vs. cuprizone treatment across 4 groups; wild-type, *Trem2*^R47H NSS^, *Trem2* ^R47H CSS^, and *Trem2* KO. **e** Heatmap of selected differentially expressed gene (DEG) from *Trem2*^R47H NSS^ (FDR<0.05 for control vs. cuprizone) compared across mouse models (see color scheme in b). **f** List of inflammation DEG upregulated in cuprizone compared to control diet that are found to be either *Trem2*-dependent (upregulated in all groups but *Trem2* KO) or *Trem2*-independent (upregulated in all groups). **g** List of uniquely upregulated DEG only found in *Trem2*^R47H NSS^. **h** List of myelination-related genes seen down- regulated in cuprizone-treated mice across all groups. **I** Representative corpus collosum 10X confocal images of wild-type, *Trem2*^R47H NSS^, *Trem2*^R47H CSS^ and *Trem2* KO on control vs cuprizone diet stained for myelin basic protein (MBP, blue), microglia (IBA1, red), and DAM gene marker (CD11c, green). **j** Quantification of IBA1^+^ cells within the field of view revealed expected increase in microgliosis in response to demyelination in cuprizone-treated mice across all groups with *Trem2* KO having significantly fewer microglia than wild-type. **k** Quantification of CD11c^+^ microglia cell volume normalized to microglia volume. n=4-8. Data are represented as mean ± SEM. Statistical significance is denoted by * p<0.05, ** p<0.01, ***p<0.001, ****p<0.0001.

Volcano plots for control vs. cuprizone across the 4 groups reveal that wild-type, *Trem2*^R47H NSS^, and *Trem2*^R47H CSS^ groups all show clear upregulation of inflammatory genes in response to cuprizone, which is markedly suppressed in the *Trem2* KO group (Fig. 1d). We further selected differentially expressed genes (DEGs) from the *Trem2*^R47H NSS^ mice (FDR<0.05 for control vs. cuprizone) and created a heatmap to compare the response to that in the other 3 groups (Fig. 1e). Upregulated genes are all associated with inflammation and are mostly shared with the wild- type and *Trem2*^R47H CSS^ groups and include disease- associated microglia (DAM) genes such as *Apoe*, *Clec7a*, and *Itgax*. Some inflammatory genes are also upregulated in *Trem2* KO mice suggesting their induction is *Trem2* independent and includes more classical inflammation related genes such as *C1qa*, *Hmox1*, *Tyrobp*, and *Trem2* itself (Fig. 1f). Downregulated genes include a unique set not altered in wild-type, *Trem2*^R47H CSS^, or *Trem2* KO groups, and are associated with dopaminergic signaling in the striatum (Fig. 1g). Shared downregulated genes are associated with myelin and oligodendrocytes, including *Cldn11*, *Mal*, *Mobp*, *Opalin*, and *Plp1* (Fig. 1h), suggesting that demyelination induced by cuprizone is not dependent upon the *Trem2*-dependent inflammation. We screened for similarities in changes in gene expression in response to cuprizone between the four groups (Supplemental Fig. 1a). Notably, *Trem2*^R47H CSS^ appears to induce a large number of DEG’s not seen in any other group (1446 genes, or 79% of all DEGs; Supplemental Fig. 1a).

To further explore gene expression changes across the groups, we performed analyses to look at functional networks of correlated genes (Weighted correlation gene network analysis (WGCNA)) and identified three modules associated with cuprizone treatment (Supplemental Fig. 1c; Cyan, Turquoise, and Royalblue modules). Turquoise module eigengene values were increased with cuprizone only in the *Trem2*^R47H NSS^ mice (Supplemental Fig. 1d), and gene ontology was associated with DNA repair and tRNA’s (Supplemental Fig. 1e). Cyan module eigengene values were increased with cuprizone to similar extents in wild-type and *Trem2*^R47H NSS^ mice, less in *Trem2*^R47H CSS^ mice, and minimally in *Trem2* KO mice (Supplemental Fig. 1f), with gene ontology being associated with inflammation (Supplemental Fig. 1g). Finally, Royalblue module eigengene values were reduced in all but the wild-type mice with cuprizone (Supplemental Fig. 1h), and gene ontology associated with myelination (Supplemental Fig. 1i), suggesting that the presence of a mutant or null *Trem2* exacerbates the demyelinating effects of cuprizone, relative to wild-type *Trem2*.

Histology for microglia (IBA1) and the DAM marker CD11c (encoded by *Itgax*) reveals extensive microgliosis in the corpus callosum of cuprizone treated wild-type, *Trem2*^R47H NSS^, and *Trem2*^R47H CSS^ mice, and to a lesser extent *Trem2* KO mice (Fig. 1i, j). While CD11c expression is induced in cuprizone treated wild-type, *Trem2*^R47H NSS^, and *Trem2*^R47H CSS^ mice, the inducation is absent in the *Trem2* KO mice (Fig. 1k), confirming the gene expression data (*Itgax*). Collectively, these results show that *Trem2*^R47H NSS^ mice show appropriate expression of *Trem2* transcripts, and do not function as a null allele as assessed by cuprizone challenge.

### *Trem2*^R47H NSS^ modulates plaque deposition in a sex-dependent fashion

To investigate the contributions of *Trem2*^R47H^ to the pathogenesis of AD, we bred *Trem2*^R47H NSS^ mice with 5xFAD mice to generate 4 groups: (i) wild-type, (ii) homozygous *Trem2*^R47H NSS^, (iii) 5xFAD, and (iv) 5xFAD; homozygous *Trem2*^R47H NSS^. We used *Trem2*^R47H NSS^ homozygotes rather than as heterozygotes that are more common in human populations, to exacerbate phenotypes associated with the R47H mutation. Hereafter, for simplicity we refer to the *Trem2*^R47H NSS^ genotype as *Trem2*^R47H^. Mice were aged to 4 and 12 mo and analyzed. Four-month old 5xFAD mice are in their rapid plaque growth stage, which then plateaus throughout the brain by ∼8-12 months depending on brain region (Forner et al., 2021). The presence of *Trem2*\*R47H induces a robust sex difference in the manifestation of Thio-S plaques in the cortex, with lower plaque density in male mice and higher in female 5xFAD/ *Trem2*^R47H^ mice vs. 5xFAD (Fig. 2c-d). Similar sex differences for plaque densities have been reported for *Trem2* KO mice crossed with 5xFAD mice (Delizannis et al., 2021). Plaque densities are similar for both 5xFAD and 5xFAD/ *Trem2*^R47H^ mice in the subiculum – a brain region that first and rapidly develops plaques in these mice. However, plaques in 5xFAD/ *Trem2*^R47H^ mice are smaller than in 5xFAD mice (Fig. 2e, f). Plotting the distribution of plaque volumes in both lines show that 5xFAD/ *Trem2*^R47H^ mice have more smaller and fewer larger plaques (Fig. 2g). Additionally, mean plaque intensity is lower in plaques from 5xFAD/*Trem2*^R47H^ mice collectively suggestive of less compaction than plaques in 5xFAD mice (Fig. 2f). Quantification of plaques at the 12-month time point show no differences in the cortex (Fig. 2j-k), but a trend towards higher plaque densities in the subiculum of 5xFAD/ *Trem2*^R47H^ mice (Fig. 2l), which are overall smaller (Fig. 2m). Consistent with plaque volume distribution from the 4-month-old animals, plaques in the subiculum of 12-month-old animals also show more smaller and fewer larger plaques in 5xFAD/ *Trem2*^R47H^ mice (Fig. 2n). However, no change in mean plaque intensity is seen suggesting that any impaired plaque compaction seen at 4 months has dissipated (Fig. 2o).

**Figure 2:**
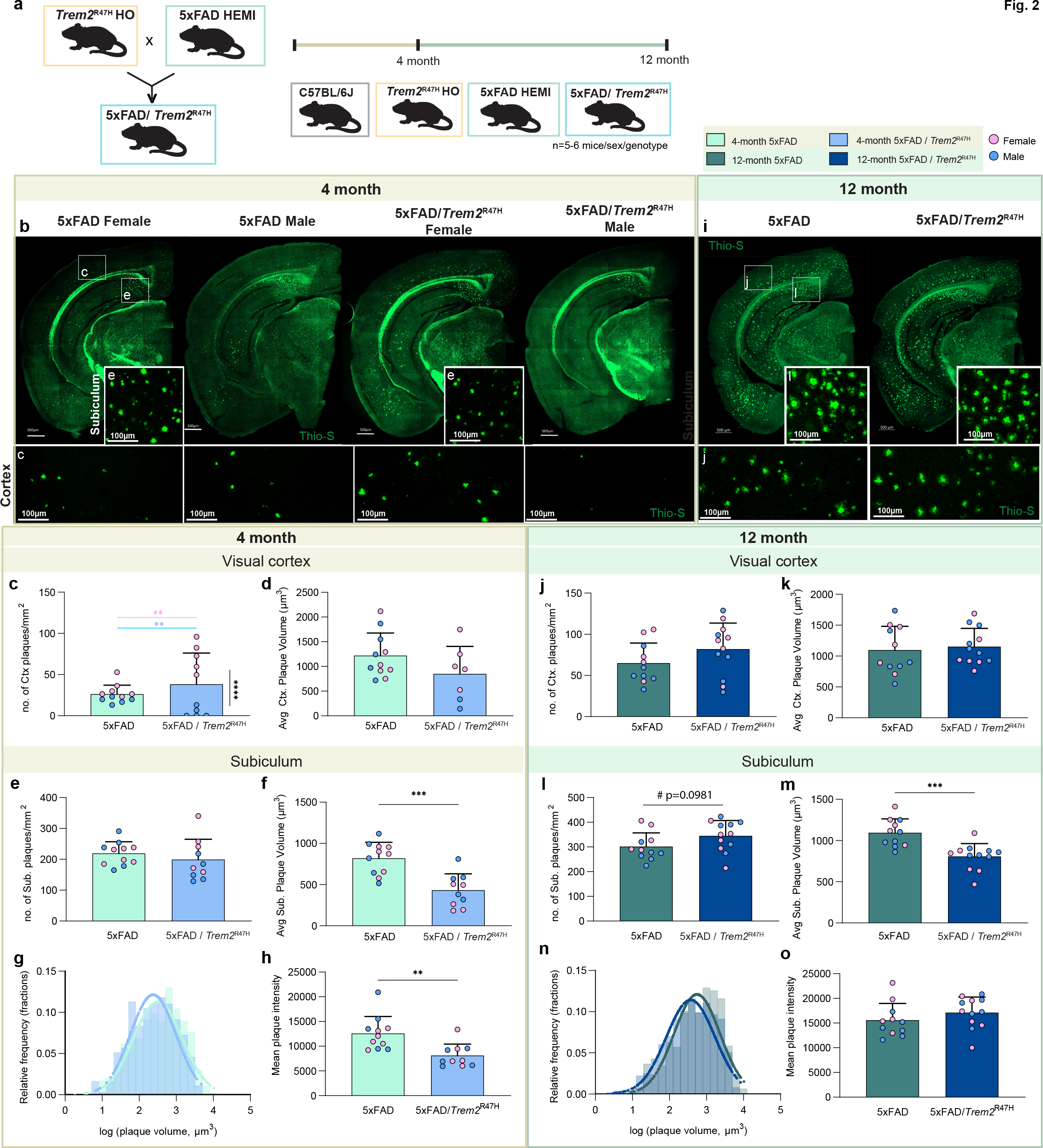
Higher amyloid plaque load in the cortex of female 5xFAD/*Trem2*^R47H^ mice. **b,i** Representative whole-brain images of 4-month-old (**b**) and 12-month-old (**i**) 5xFAD and 5xFAD/ *Trem2*^R47H^ stained for dense-core plaques using Thioflavin-S (green) with insets of 20X confocal images of the visual cortex (b) and subiculum (d). Scale bar = 500µm. **c** At 4-month, quantification of Thio-S^+^ density reveals significant sex difference in dense-core plaque burden within 5xFAD/ *Trem2*^R47H^ in the visual cortex (unpaired t-test, p<0.0001). Female 5xFAD/*Trem2*^R47H^ also exhibited heavier plaque burden in the visual cortex compared to the age-match 5xFAD (2-way ANOVA, p<0.001). **d, f** Average plaque volumes showed no difference in the visual cortex (**d**) but significantly decreased in the subiculum of 5xFAD/ *Trem2^R47H^* compared to 5xFAD (**f**, p<0.001). **j, l** At 12-month, quantification of dense-core plaque numbers per mm^2^ revealed no significant difference between 5xFAD and 5xFAD/ *Trem2*^R47H^ in the visual cortex (**j**) but a trending increase in the subiculum (**l,** p=0.0981). **k, m** No significant difference was observed in average plaque size in the cortex (**j**) but plaque size decreased in 5xFAD/ *Trem2^R47H^* vs 5xFAD in the subiculum (**m,** unpaired t-test, p<0.001). **g, n** Plaque volume relative frequency histogram superimposed by the best fit of data on a Gaussian distribution curve showed a shift in plaque size between 5xFAD and 5xFAD/ *Trem2*^R47H^ at 4-months (**g**) and 12-months (**n**). **h, o** Total mean plaque intensity calculation revealed lower intensity in 5xFAD/*Trem2*^R47H^ compared to 5xFAD at 4-months (**h**) but not at 12-months (**o**). n=10-12. Data are represented as mean ± SEM. Statistical significance is denoted by * p<0.05, ** p<0.01, ***p<0.001, ****p<0.0001. Statistical trends are given by # 0.05<p<0.1.

Aβ40 and Aβ42 were measured in detergent soluble and insoluble fractions from microdissected hippocampi and cortices. At 4 months of age, increased Aβ40 and 42 are found in the 5xFAD/ *Trem2*^R47H^ hippocampus and a trending reduction in Aβ42 in the cortex, but no difference in the insoluble fraction in either brain region (Fig. 3a-h). By 12 months of age Aβ levels are greatly increased in both brain regions and fractions, with increased soluble Aβ42 levels in the hippocampus and increased insoluble Aβ40 and 42 in the cortex of 5xFAD/ *Trem2*^R47H^ mice (Fig. 3i-p). Collectively, these results show that *Trem2*\*R47H impacts the level of both plaque and Aβ in a brain region and age-specific manner.

**Figure 3:**
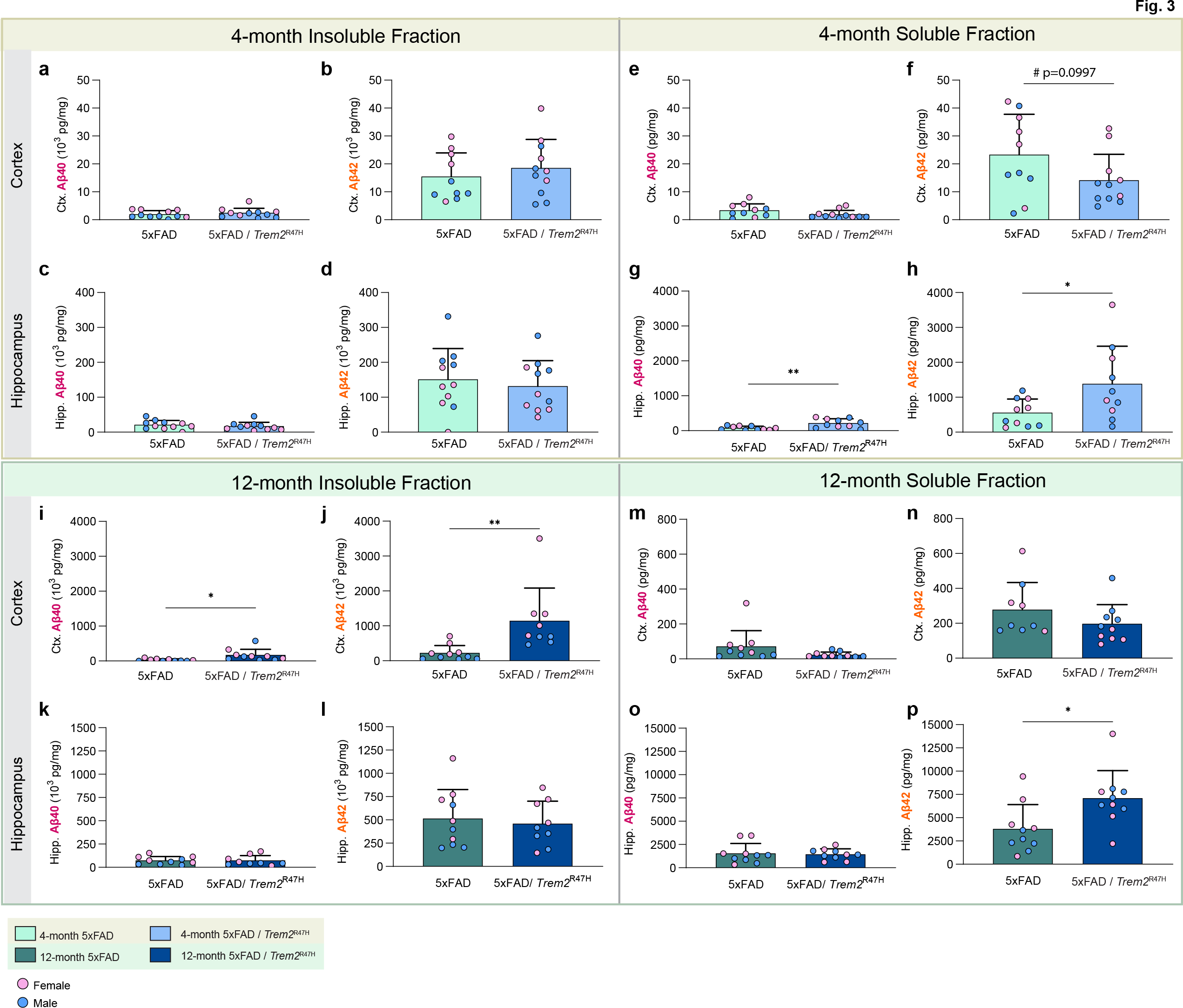
Quantification of insoluble and soluble Aß in micro-dissected hippocampi and cortices using Meso Scale Discovery technology. **a-d** Quantification of 4-month cortical and hippocampal insoluble fraction revealed no difference in Aβ40 and Aβ42 in 5xFAD/ *Trem2^R47H^* vs 5xFAD. e-h In the 4-month soluble fraction, there is no difference in cortical Aβ40 (e), but a trending decrease in Aβ42 level was observed in the cortical fraction of 5xFAD/ *Trem2^R47H^* vs 5xFAD (f, p=0.0997). Increases in Aβ40 (g, p<0.05) and Aβ42 levels (h, p=0.01) were observed in hippocampal fraction of 5xFAD/ *Trem2^R47H^* compared to 5xFAD. i-l At 12-month, insoluble cortical Aβ40 (i, p<0.05) and Aβ42 (j, p<0.01) are increased in 5xFAD/ *Trem2*^R47H^ compare to 5xFAD while no difference was observed in hippocampal fraction (k, l). m-p No difference was observed in soluble fraction except for an increase in hippocampal Aβ42 in 5xFAD/ *Trem2*^R47H^ (p, p<0.05). Data are represented as mean ± SEM. Statistical significance is denoted by * p<0.05, ** p<0.01, ***p<0.001, ****p<0.0001. Statistical trends are given by # 0.05<p<0.1.

### Initial impairment in microglia-plaque interactions is lost with age/disease progression

Given the expression of TREM2 by microglia in the brain, we explored microglial densities and morphologies. Morphological analyses of non-plaque, cortical IBA1^+^ microglia showed increased process length/cell but decreased diameter in *Trem2*^R47H^ compared to WT mice (Fig. 4c, d). At 4- months, as expected, the sex difference observed in plaque load of 5xFAD/*Trem2*^R47H^ is reflected in cortical microglia density (Fig. 4e). Interestingly, while microglia density is unchanged, average cortical microglia volume decreases significantly in 5xFAD/ *Trem2*^R47H^ compared to 5xFAD (Fig. 4f). In the subiculum, where Thio-S^+^ plaques are abundant, IBA1^+^ microglia density and size are increased in 5xFAD and 5xFAD/ *Trem2*^R47H^ compared to wild-type mice. Moreover, IBA1^+^ cell densities and volumes of 5xFAD/ *Trem2*^R47H^ remain lower than 5xFAD (Fig. 4g, h). Notably, in both brain regions, *Trem2*^R47H^ mice exhibit lower IBA1^+^ cell volume compared to WT, indicating that the *Trem2*\*R47H variant elicits a plaque-independent effect on microglia morphology (Fig. 4f, h). The reduction in volume of IBA1^+^ cells occurs in tandem with a lack of plaque-microglia interaction in 5xFAD/ *Trem2*^R47H^ shown through quantification of Thio-S and IBA1 colocalization in the subiculum with a further impairment in female 5xFAD/ *Trem2*^R47H^ mice compared to males (Fig. 4o). Notably, the decreased microglia volume in *Trem2*^R47H^ mice and 5xFAD/ *Trem2*^R47H^ compared to WT and 5xFAD, respectively, is absent at 12-month (Fig. 4f, h, j, l). Moreover, the lack of plaque-microglia interaction, along with the sex-difference in 5xFAD/ *Trem2*^R47H^ at 4- months is absent at 12-months in the subiculum, suggesting age/disease-dependent changes in the *Trem2*\*R47H variant effect on microglia morphology and function (Fig. 4o, p).

**Figure 4:**
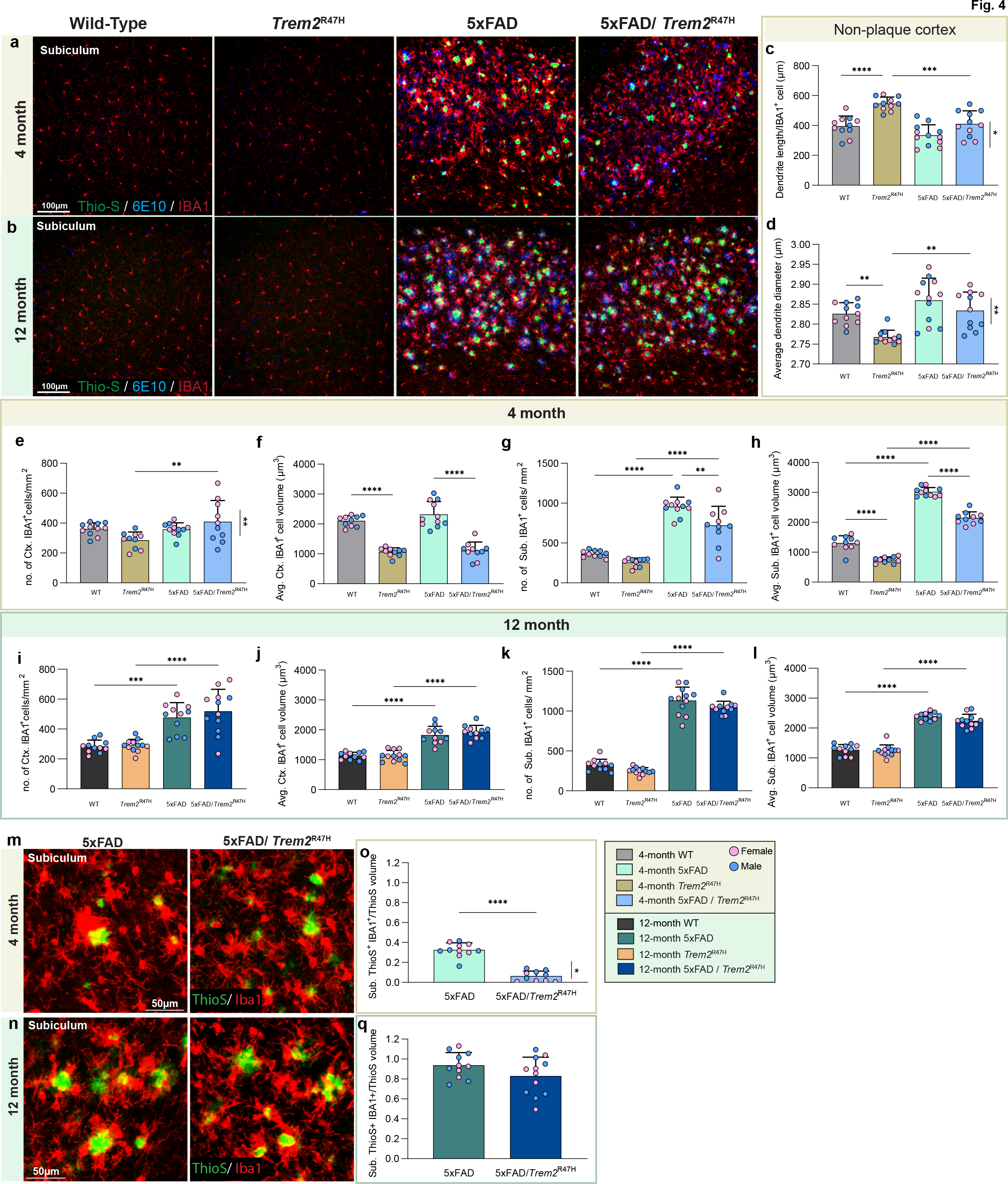
Age/disease-dependent impairment of plaque-microglia interaction driven by *Trem2*\*R47H. **a, b** Subiculum - representative confocal images of wild-type, *Trem2*^R47H^, 5xFAD, and 5xFAD/ *Trem2*^R47H^ at 4- (**a**) and 12-month (**b**) stained with Thio-S for dense-core plaques (green), immunolabeled with 6E10 for diffused plaque (blue), and IBA1 for microglia (red). **c, d** Quantification of cortical microglia morphology in non-plaque regions revealed increased dendrite length per IBA1^+^ cell in *Trem2*^R47H^ compared to WT and 5xFAD/ *Trem2*^R47H^ (**c**) but decreased average dendrite diameter (**d**). **e- h** Quantification of IBA1^+^ cell density and average volume in the visual cortex (**e, f**) and subiculum (**g, h**) at 4-month of age**. e, f** In the cortex, a sex-dependent decrease in microglia number (5xFAD/ *Trem2*^R47H^ female vs male: **e**, p<0.01) and a decrease in average microglial volume in the presence of *Trem2*^R47H^ (**f**, WT vs *Trem2*^R47H^ and 5xFAD vs 5xFAD/ *Trem2*^R47H^, p<0.0001) were found. **g, h** In the subiculum, quantification revealed decreased in 5xFAD/ *Trem2*^R47H^ compared to 5xFAD (**g**, cell density, p<0.01; **h**, cell volume, p<0.0001). **i -l** Quantification of IBA1^+^ cell density and average volume in the visual cortex (**i, j**) and subiculum (**k, l**) at 12-month-old. **m-n** Representative 20x images of Thio-S (green) and IBA1 (red) colocalization in the subiculum at 4-month (**m**) and 12-month (**n**). **o, q** Quantification of percent colocalized volume of Thio-S^+^ and IBA1^+^ cell normalized to total Thio-S volume per field of view in the subiculum revealed decreased plaque-microglia interaction in 5xFAD/ *Trem2*^R47H^ at 4-month with sex-differences (**o**, p<0.0001; females vs males. P<0.05) but not 12-month (**q**). n=10-12. Data are represented as mean ± SEM. Statistical significance is denoted by * p<0.05, ** p<0.01, ***p<0.001, ****p<0.0001.

### *Trem2*\*R47H produces increased brain damage in response to plaques

Microglia form a protective barrier around plaques, while contributing to their compaction and growth (Condello et al., 2015). Given the initial impairment in microglia-plaque interactions in 5xFAD/ *Trem2*^R47H^ mice we next explored the halo of dystrophic neurites that develops around dense core plaques (Gowrishankar et al., 2015; Sadleir et al., 2016) which can be visualized with lysosomal-associated membrane protein 1 (LAMP1). Representative stains are shown (Fig. 5a, 5b). Normalization of LAMP1 volume to plaque volume reveals an increase in dystrophic neurites per plaque area in 5xFAD/ *Trem2*^R47H^ mice at 4 months of age (Fig. 5c). Consistent with the restoration of microglia-plaque interactions by 12 months of age, no difference in dystrophic neurites per plaque is seen between 5xFAD/ *Trem2*^R47H^ and 5xFAD mice at this age (Fig. 5d). 5xFAD females have more dystrophic neurites than male. Interestingly, LAMP1^+^ dystrophic neurites exhibit more dissipated morphology at 12-month compared to 4-month, consistent with disrupted axonal transport at later age/disease stages (Sharoar et al., 2021).

**Figure 5:**
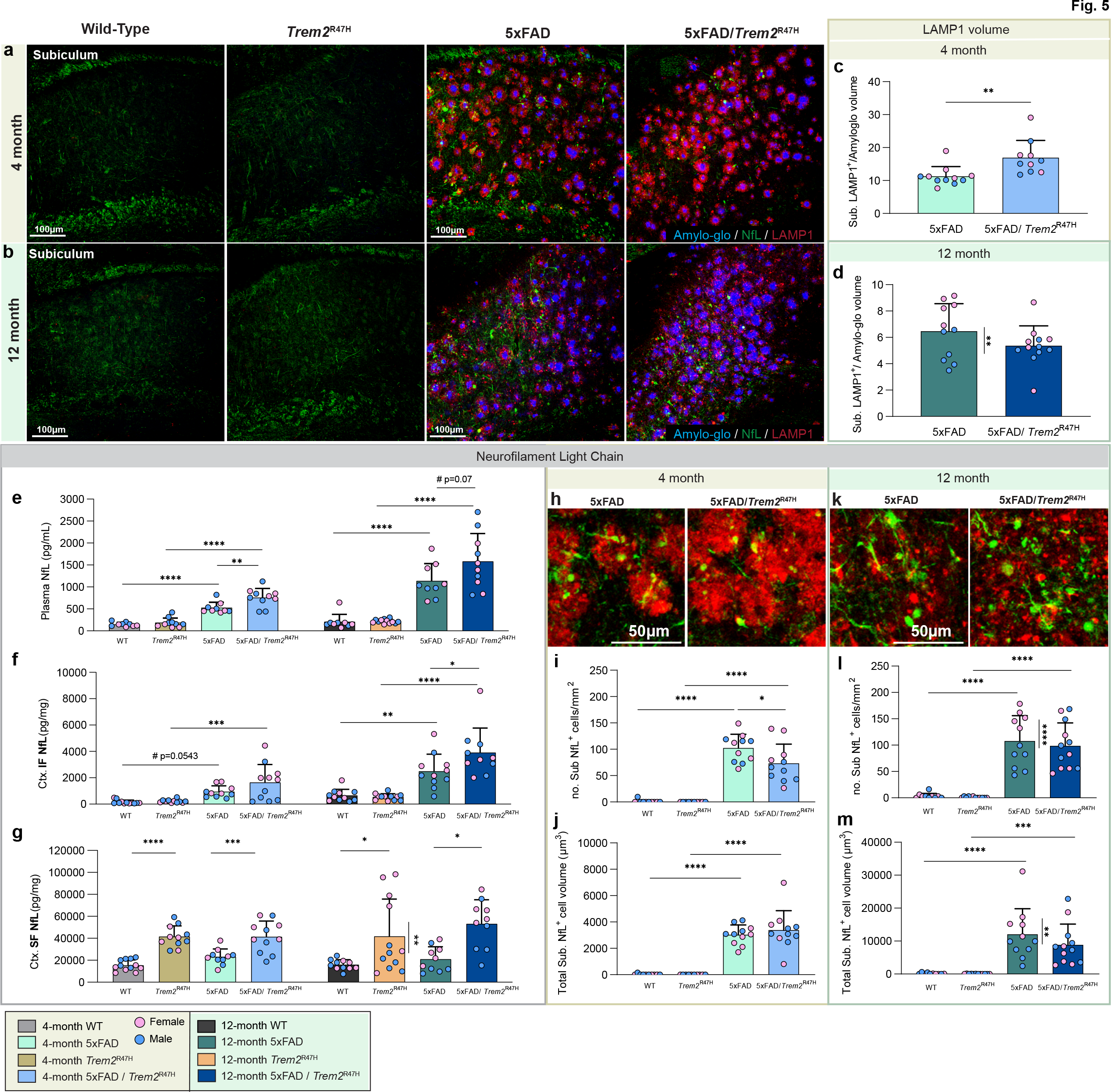
*Trem2\**R47H induces age/disease-dependent dystrophic neurites and axonal damage. **a, b** Representative 20X confocal images of wild-type, *Trem2*^R47H^, 5xFAD, and 5xFAD/ *Trem2*^R47H^ at 4- (**a**) and 12-month (**b**) stained with Amylo-Glo for dense-core plaques (blue), immunolabeled with NfL for neurofilament light chain (green), and LAMP1 for dystrophic neurites (red). **c, d** Quantification of subiculum LAMP1 volume normalized to Amylo-Glo volume revealed an increase in dystrophic neurites at 4-month (**c**, p<0.01) but not at 12-month (**d**) with sex- difference in 5xFAD highlighted (5xFAD female vs male, t-test, p<0.01). **e- g** Measurement of NfL in plasma (**e**), cortical insoluble fraction (**f**), and soluble fraction (**g**) via Meso Scale Discovery technology revealed consistent increase in NfL level in 5xFAD/ *Trem2*^R47H^ at both 4- and 12- month. **h, k** Representative higher magnification images of immunolabeled NfL spheroids (green) colocalized with LAMP1 (red) in the subiculum of 4-month (**h**) and 12-month (**k**) 5xFAD and 5xFAD/ *Trem2*^R47H^. **i, j** Quantification of NfL^+^ spheroids number per mm^2^ showed a decrease in 5xFAD/ *Trem2*^R47H^ compared to 5xFAD (**i**, p<0.05) but no change in spheroid volume (**j**) at 4- month. **l, m** Quantification revealed no change in both number (**l**) or volume (**m**) of NfL^+^ spheroids between 5xFAD and 5xFAD/ *Trem2*^R47H^ but revealed a sex-difference in 5xFAD at 12-Month. n=10-12. Data are represented as mean ± SEM. Statistical significance is denoted by * p<0.05, ** p<0.01, ***p<0.001, ****p<0.0001.

Neurofilament light chain (NfL) is emerging as a clinically useful plasma biomarker for damage occurring in the brain, including in AD where it tracks with cortical thinning and cognitive decline (de Wolf et al., 2020; Lee et al., 2022; Quiroz et al., 2020), while in mouse models of AD it correlates with plaque load (Javonillo et al., 2021). We measured plasma NfL as a surrogate marker of brain damage and found it increased in 5xFAD mice compared to wild-type mice at 4 months of age, which increases further at 12, consistent with plaque load (Fig. 5e). The presence of *Trem2*\*R47H further increased plasma NfL in 5xFAD at both ages (Fig. 5e), consistent with the hypothesis that TREM2 dysfunction exacerbates brain damage. To explore the transition of brain NfL to the plasma we measured NfL levels in the detergent soluble and insoluble fractions of microdissected cortices. NfL in the soluble fraction is not increased in 5xFAD mice at either 4 or 12 months of age, but soluble NfL is increased by the presence of *Trem2*\*R47H in either wild- type or 5xFAD mice (Fig. 5f). However, NfL in the insoluble fraction aligns with levels in the plasma, with increases seen in 5xFAD compared to wild-type mice at 4 months, with further increases at 12 (Fig. 5g). As with plasma, levels further trend higher in 5xFAD/ *Trem2*^R47H^ mice. Thus, the presence of *Trem2*\*R47H induces changes in NfL, and further exacerbates plaque- induced increases in both detergent insoluble and plasma NfL. To identify the cellular source of NfL, we immunostained for dystrophic neurites (LAMP1) and NfL. Large spherical structures of NfL are seen in the vicinity of plaques and are absent from wild-type and *Trem2*^R47H^ mice, where staining is observed only in axonal fibers (Fig. 5a, b). Similar bead-like NfL^+^ spheroids were reported in ischemia-affected human and mouse tissues as sign of axonal damage (Mages et al., 2018). Notably, these NfL spheroids colocalize with dystrophic neurites associated with plaques (Fig. 5h, k). Quantification of NfL^+^ structures showed a decreased spheroid number in 5xFAD/ *Trem2*^R47H^ compared to 5xFAD, while having similar size at 4-month (Fig. 5i, j). At 12-month, no difference in disrupted/extracellular NfL number or size is observed between 5xFAD and 5xFAD/ *Trem2*^R47H^. However, in 5xFAD, there is a sex difference with females having more NfL compared to males, which is also observed in LAMP1^+^ dystrophic neurite amount (Fig. 5l, m). Collectively, these findings reveal associations between plaques and dystrophic neurites with NfL accumulation and its transition to the insoluble fraction and plasma NfL.

### *Trem2*^R47H^ initially protects against plaque induced LTP and synaptic deficits

With increased dystrophic neurites induced by the R47H variant, we investigated short- and long- term synaptic plasticity in WT, *Trem2*^R47H^, 5xFAD, and 5xFAD/*Trem2*^R47H^ hippocampi via theta burst-induced pair-purse facilitation (PPF) and long-term potentiation (LTP) in acute hippocampal slices. Consistent with our previous findings (Forner et al., 2021), 5xFAD have impaired LTP from 4 months of age. Remarkably, this impairment is supressed by the presence of *Trem2*^R47H^ (Fig. 6a, b), while no change in PPF is observed (Fig. 6c). Consistent with a lack of LTP impairment in 5xFAD/*Trem2*^R47H^ mice, immunostaining of pre- and post-synaptic elements for synaptophysin and PSD-95, respectively, revealed a decrease in pre-synaptic puncta in 5xFAD animals, which is prevented in 5xFAD/*Trem2*^R47H^ mice (Fig. 6d,e), and no changes in post-synaptic elements across all groups (Fig. 6f). In contrast, by 12-months of age, hippocampal LTP deficits are seen in *Trem2*^R47H^, 5xFAD, and 5xFAD/*Trem2*^R47H^ hippocampi compared to WT animals (Fig. 6g, h). Notably, PPF responses at 12-months show significant decreases in presynaptic plasticity in 5xFAD and 5xFAD/ *Trem2*^R47H^ compared to their wild-type controls at 40ms stimulus interval, however, the effect in 5xFAD/ *Trem2*^R47H^ is absent at 100ms interval (Fig. 6i). Staining of presynaptic puncta also reflects this deficit in 5xFAD and 5xFAD/ *Trem2*^R47H^ while also capturing the decrease in Synaptophysin^+^ staining of *Trem2*^R47H^ compared to WT, as well as 5xFAD/ *Trem2*^R47H^ vs 5xFAD (Fig. 6l). Postsynaptic elements are also decreased in *Trem2*^R47H^ compared to WT hippocampi, along with an increase in 5xFAD/ *Trem2*^R47H^ vs 5xFAD, which reflect the observed fEPSP and mean potentiation data (Fig. 6m).

**Figure 6:**
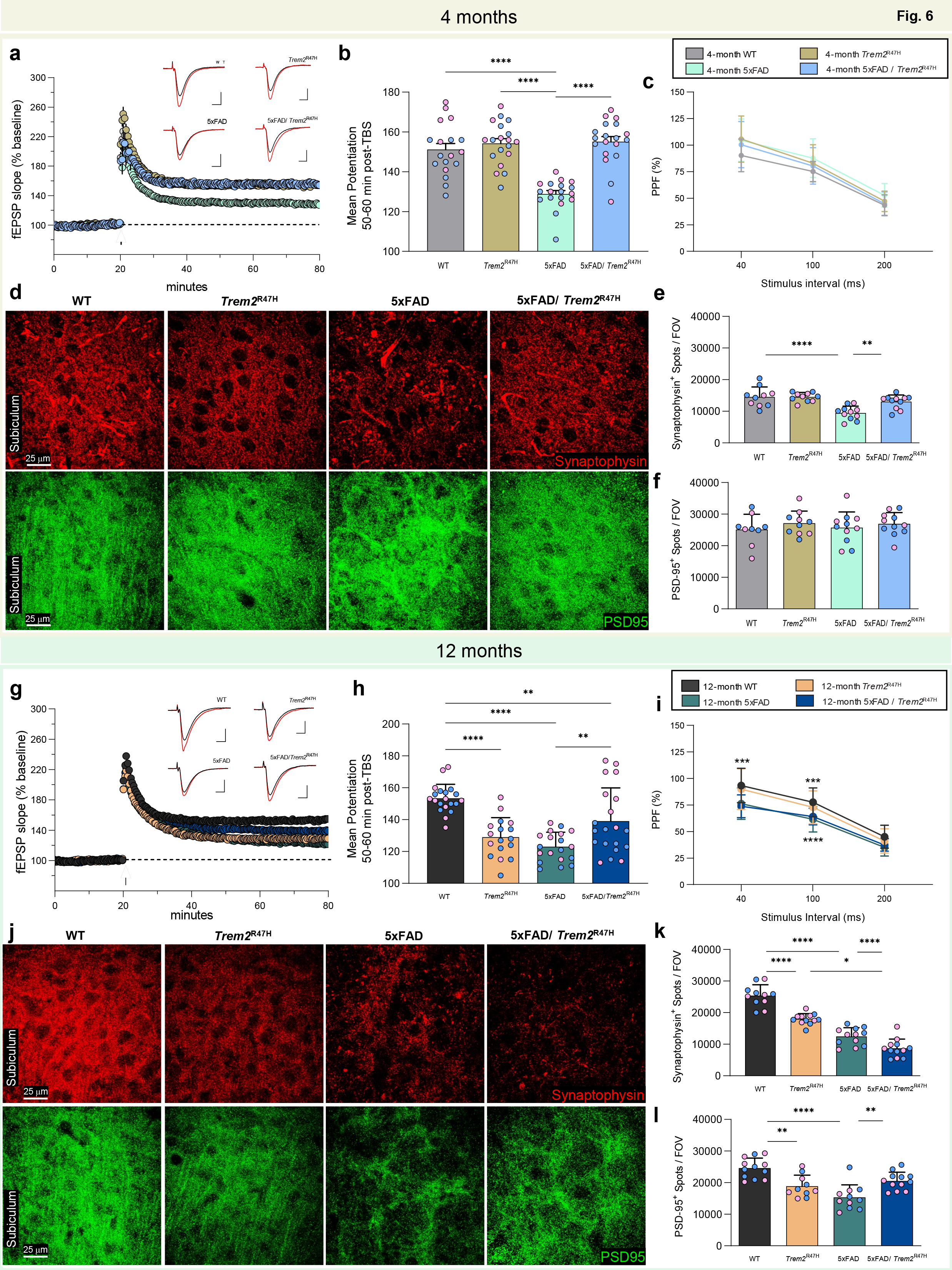
Age-dependent LTP deficit. Hippocampal slices of 4 and 12-month-old WT, *Trem2*^R47H^, 5xFAD, and 5xFAD/ *Trem2*^R47H^ mice of both sexes were analyzed using theta-burst induced long-term potentiation (LTP). **a** Time course of fEPSP slope (as percentage of baseline) following theta burst stimulation (TBS, black arrow) of 4-month WT, *Trem2*^R47H^, 5xFAD, and 5xFAD/ *Trem2*^R47H^ mice showing impaired LTP in 5xFAD but not 5xFAD/ *Trem2*^R47H^. Insets show field synaptic responses collected during baseline (black line) and 1 hr after TBS (red line). Scale: 1mV/5ms. **b** Mean potentiation (±SEM) during the last 10 min of recording in slices from 4 months WT, *Trem2*^R47H^, 5xFAD, and 5xFAD/ *Trem2*^R47H^ mice shows reduction in 5xFAD but not 5xFAD/ *Trem2*^R47H^ mice (p< 0.0001). **c** Paired-pulse facilitation (PPF) was measured at 40, 100, and 200ms intervals. At 4 months of age, no significant difference was observed between groups at any of the three intervals tested (p=0.144). **d** Representative 63x confocal images of 4-months WT, *Trem2*^R47H^, 5xFAD, and 5xFAD/ *Trem2*^R47H^ subiculum immunolabeled with synaptophysin for presynaptic elements (top panel, red) and PSD-95 for postsynaptic elements (bottom panel, green). **e** Quantification of synaptophysin^+^ spots per field-of-view (FOV) showed a decrease in presynaptic elements in 5xFAD (WT vs 5xFAD, p<0.0001; 5xFAD vs 5xFAD/ *Trem2*^R47H^; p<0.01). **f** PSD-95^+^ spots per FOV revealed no difference in postsynaptic elements. **g** Time course of fEPSP slope following theta burst (black arrow) of 12-months WT, *Trem2*^R47H^, 5xFAD, and 5xFAD/ *Trem2*^R47H^ mice show impaired LTP in *Trem2*^R47H^, 5xFAD and partial impairment in 5xFAD/ *Trem2*^R47H^. Insets show field synaptic responses collected during baseline (black line) and 1 hr after theta burst stimulation (red line). Scale: 1mV/5ms. **h** Mean potentiation (±SEM) during the last 10 min of recording in slices from 12 months WT, *Trem2*^R47H^, 5xFAD, and 5xFAD/ *Trem2*^R47H^ mice shows reduction in *Trem2*^R47H^, 5xFAD, 5xFAD/ *Trem2*^R47H^ but a partial rescue in 5xFAD/ *Trem2*^R47H^ compared to 5xFAD (WT vs *Trem2*^R47H^, 5xFAD, 5xFAD/ *Trem2*^R47H^; p< 0.0001, p< 0.0001, p< 0.01 respectively; 5xFAD vs 5xFAD/ *Trem2*^R47H^; p<0.01). **i** Significant group effect was found in PPF at 40 (WT vs 5xFAD and WT vs 5xFAD/ *Trem2*^R47H^; p=0.0002, p<0.0001 respectively) and 100 ms stimulus intervals (WT vs 5xFAD and WT vs 5xFAD/ *Trem2*^R47H^; p=0.0005, p=0.0046 respectively). **k** Representative 63x confocal images of 12-months WT, *Trem2*^R47H^, 5xFAD, and 5xFAD/ *Trem2*^R47H^ subiculum for synaptophysin (red) and PSD-95 (green). **l** Synaptophysin^+^ cells quantification showed decrease in presynaptic elements in 5xFAD compared to WT and further decrease in presence of *Trem2*\*R47H (WT vs 5xFAD, p<0.0001; WT vs *Trem2*^R47H^, p<0.0001; 5xFAD vs 5xFAD/ *Trem2*^R47H^, p<0.0001). **m** PSD95^+^ cell quantification showed decrease in postsynaptic elements in 5xFAD compared to WT and further decrease in presence of *Trem2*\*R47H (WT vs 5xFAD, p<0.0001; WT vs *Trem2*^R47H^, p<0.01; 5xFAD vs 5xFAD/ *Trem2*^R47H^, p<0.01). n=10-12. Data are represented as mean ± SEM. Statistical significance is denoted by * p<0.05, ** p<0.01, ***p<0.001, ****p<0.0001.

### *Trem2*\*R47H initially suppresses but then enhances neuroinflammation with age/disease progression, including production of a unique interferon signature

To assess gene expression changes with age and genotype we performed RNA-seq from microdissected hippocampi from 4- and 12-month-old WT, *Trem2*^R47H^, 5xFAD, and 5xFAD/ *Trem2*^R47H^ mice. Volcano plots illustrate the global changes in gene expression between *Trem2*^R47H^ and WT mice, 5xFAD and WT, and 5xFAD/ *Trem2*^R47H^ and 5xFAD mice at both 4 and 12 months (Fig. 7a). Gene expression data can be explored in an interactive fashion at http://swaruplab.bio.uci.edu:3838/5xFAD_Trem2/. Many genes are significantly altered in *Trem2*^R47H^ hippocampi compared to WT, with a group of downregulated genes manifesting at 12 months. Differentially expressed genes (DEGs) in 5xFAD mice vs. WT mice at both ages are mostly upregulated genes and represent the strong inflammatory response seen in these mice, and include DAM genes such as *Cst7*, *Itgax*, *Clec7a*, as well as *Trem2*. DEGs between 5xFAD/ *Trem2*^R47H^ and 5xFAD mice at 12 months are plotted as a heatmap (Fig. 7b; FDR < 0.05, no FC cutoff). Many of these DEGs represent downregulated genes that appear in *Trem2*^R47H^ mice both on a WT and 5xFAD background and are considered further below (Fig. 7c – Royalblue module). The 13 genes that are changed due to the interaction between 5xFAD and 5xFAD/ *Trem2*^R47H^ genotypes are bolded (Fig. 7b). Notably, this subset consists of inflammation related genes, and show downregulation of *Itgax*, *Tgfb1*, and *Ccl6* compared to 5xFAD mice, but an upregulation of many genes associated with interferon signaling, such as *Ifi47*, *Ifit1-3*, and *Gbp2*, *6*, and *7*.

**Figure 7:**
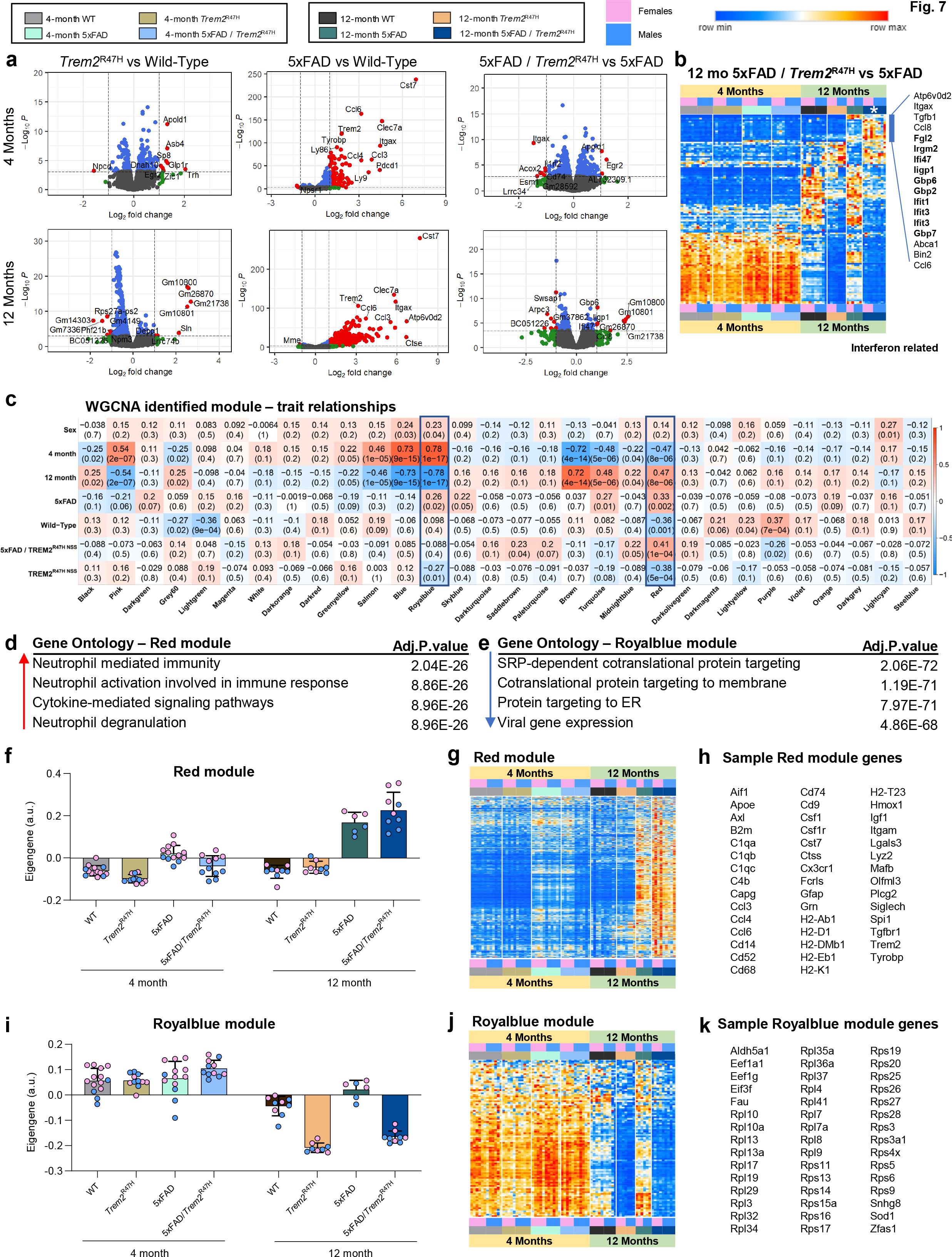
*Trem2*\*R47H initially suppresses but then enhances neuroinflammation with age/disease 35rogression. **a** Volcano plot of DEGs, displaying fold change of gene expression (log2 scale) and *P* values (−log10 scale) at 4- and 12-months between wild-type vs *Trem2*^R47H^, 5xFAD vs wild-type, and 5xFAD/ *Trem2*^R47H^ vs 5xFAD. **B** Heatmap generated from selected DEG between 5xFAD and 5xFAD/ *Trem2*^R47H^ at 12-months, showing the list of uniquely upregulated genes only in 5xFAD/ *Trem2*^R47H^ with interferon-related genes in bold. **C** module Trait relationship heatmap from WGCNA to explore the relationship between sex, genotype, and age. The color is based on their correlation: red is a strong positive correlation, while blue is a strong negative correlation. Royalbule and red modules were chosen to investigate more based on their correlation across all data trait. **D** Gene oncology of the red module revealed increased in neutrophil responses and pathways. **E** Royalblue module gene oncology is related to protein translation. **F** Eigengene of the red module plotted as bar graphs for WT, *Trem2*^R47H^, 5xFAD and 5xFAD/ *Trem2*^R47H^ at 4- and 12-month. **g** Heatmap generated from selected DEG in the red module. **H** List of sample genes from the red modules. **I** Eigengene of the Royalblue module plotted as bar graphs for WT, *Trem2*^R47H^, 5xFAD and 5xFAD/ *Trem2*^R47H^ at 4- and 12-month **j** Heatmap generated from selected DEG in the Royalblue module. **H** List of sample genes from the Royalblue modules.

To further explore gene expression changes across the groups we performed analyses to look at functional networks of correlated genes (WGCNA). We identified one module associated with all genotypes (Red), and one module associated with *Trem2*^R47H^ genotype (Royalblue; Fig. 7c). Red module gene ontology was associated with inflammation and immune related genes (Fig. 7d), while Royalblue gene ontology was associated with protein translation (Fig. 7e). Plotting of eigengene values for the Red module for all groups (Fig. 7f), revealed an increase in 4 month old 5xFAD mice compared to both WT and *Trem2*^R47H^ mice, corresponding to the inflammatory response to the plaques in those mice. Notably, no such increase in eigengene value is observed in 4-month-old 5xFAD/ *Trem2*^R47H^ mice, showing a suppression of inflammation at the 4-month timepoint, particularly in male mice, mirroring the LTP and synaptic deficits seen in these mice. By 12 months, however, robust increases in Red module eigengene values are seen in both 5xFAD and 5xFAD/ *Trem2*^R47H^ mice, showing that the initial suppression of inflammation induced by the *Trem2*^R47H^ variant dissipates with time/age/disease progression, in line with the reults of histology. A heatmap of Red module genes is shown in (Fig. 7g), and sample genes in Fig. 7h, including *Apoe*, *Axl*, *C1qa-c*, *Cd74*, *Csf1r*, *Cst7*, *Grn*, *Igf1*, and *Trem2* itself. Similarly, we plotted eigengene values for the Royalblue module, representing genes that are downregulated with *Trem2*^R47H^ genotype at 12 months of age (Fig. 7i), along with a heatmap of the module genes (Fig. 7j), and samples genes (Fig. 7k), revealing strong representation of ribosome associated genes.

Given the selective downregulation of *Itgax* at both 4- and 12-months-of-age between 5xFAD/ *Trem2*^R47H^ and 5xFAD mice (Fig. 8c, g), and *CD74* only at 4 months of age (Fig. 8d, h), we performed immunohistochemistry for both markers (Fig. 8e, f, I, j), alongside microglia and plaques. CD11c (*Itgax*) and CD74 staining were seen in a subset of plaque-associated microglia, with greater numbers seen at 12- vs. 4-months of age. Concordant with gene expression data, reduced CD11c staining was observed in 5xFAD/ *Trem2*^R47H^ mice at both timepoints, while CD74 staining is only reduced at 4-months of age (Fig. 8e, f, I, j).

**Figure 8:**
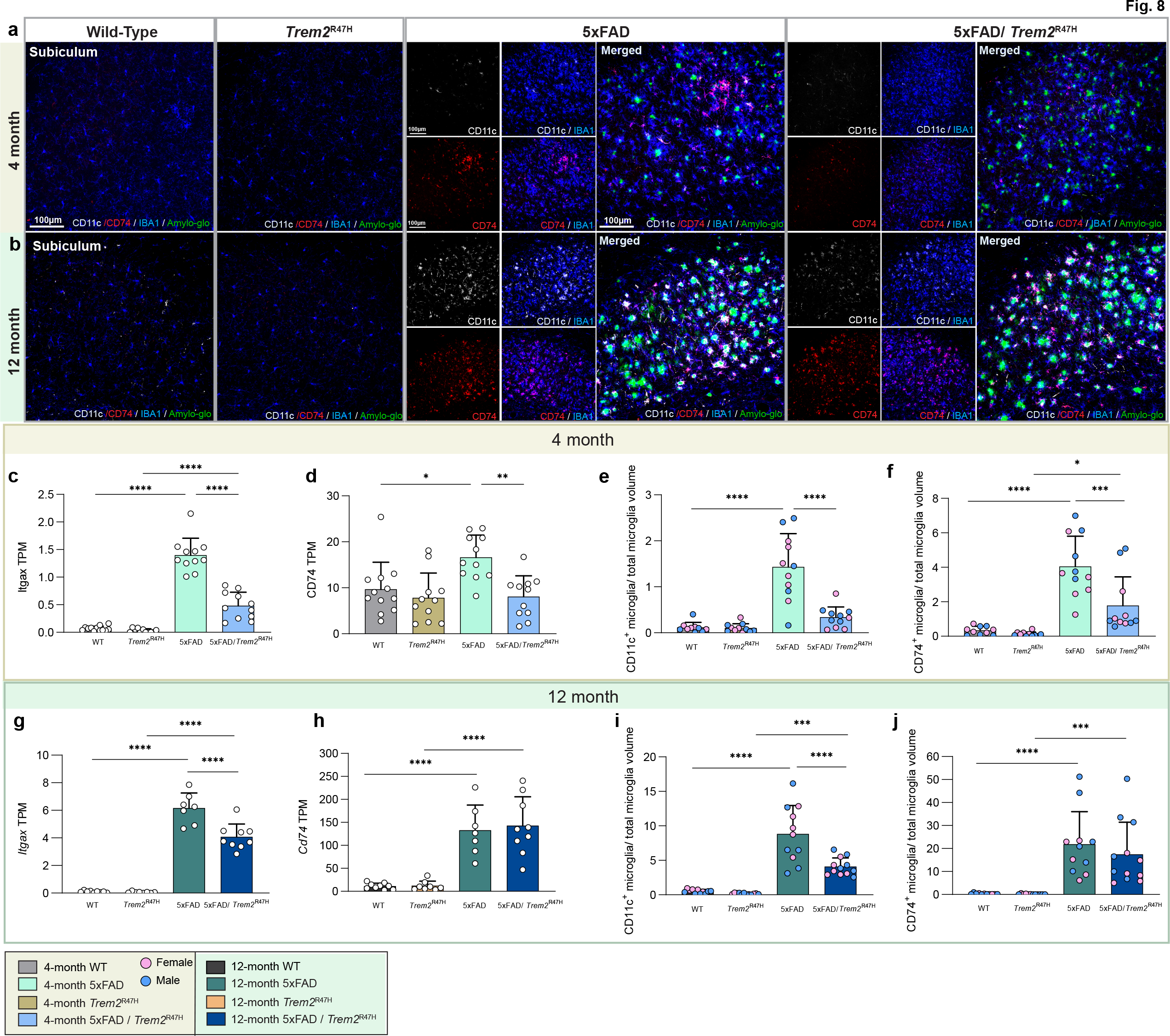
*Trem2*\*R47H reduces expression of *itgax and Cd74*. **a, b** Representative confocal images of hippocampal subiculum wild-type, *Trem2*^R47H^, 5xFAD, and 5xFAD/ *Trem2*^R47H^ at 4- (**a**) and 12-month (**b**) stained with Amylo-Glo for dense-core plaques (green), immunolabeled with IBA1 for microglia (blue), CD11c (white), and CD74 (red) confirms reduced expression of *itgax* and *Cd74*. **c, d** 4-month TPM values of *itgax* (CD11C) and *Cd74* plotted as bar graphs showed decreased expression between 5xFAD and 5xFAD/ *Trem2*^R47H^ in both *itgax* (**b**, p<0.0001) and *Cd74* (**c**, p<0.01) **e, f** At 4-month, quantification of colocalization of IBA1 and CD11c (**e**) and CD74 (**f**) normalized to total IBA1 volume confirmed reduction in the respective gene expression (5xFAD vs 5xFAD/ *Trem2*^R47H^; **e** p<0.0001, **f** p<0.001). **g, h** 12-month TPM values of *itgax* and *Cd74* from bulk cortical RNA sequencing data revealed decreased in *itgax* TPM value (**g**, p<0.0001) but not *Cd74*. **i, l** Quantification of 12-month colocalization of IBA1 and CD11c (**i**) and CD74 (**l**) normalized to total IBA1 volume confirmed changes in the respective gene expression (5xFAD vs 5xFAD/ *Trem2*^R47H^; **i** p<0.0001). n=10-12. Data are represented as mean ± SEM. Statistical significance is denoted by * p<0.05, ** p<0.01, ***p<0.001, ****p<0.0001.

Collectively, these results show that the *Trem2*^R47H^ variant induces age-related changes in genes involved in translation, but also mediates an initial suppression of inflammation in response to plaques in 5xFAD mice, which dissipates then becomes exacerbated with age/disease progression. Furthermore, the *Trem2*^R47H^ variant generates a unique interferon-related gene expression signature in 5xFAD mice.

## DISCUSSION

The identification of coding sequence changes in *TREM2* that were strongly associated with increased risk for development of LOAD focused interest on both TREM2 function and microglia that predominantly express TREM2 in the brain (Guerreiro et al., 2013; Jonsson et al., 2013). Initial studies of *TREM2* KO mice found that TREM2 was necessary for the microglial reaction to plaques, as well as their transition to a “DAM” phenotype, characterized by the specific expression of genes such as *Cst7, Clec7a, Itgax*, and *Apoe* (Keren-Shaul et al., 2017). Furthermore, the absence of TREM2, coinciding with a lack of microglial reaction to plaques, appeared to paradoxically worsen disease progression (Meilandt et al., 2020; Wang et al., 2016; Yuan et al., 2016a). Although these KO studies validated TREM2 as a key and central player for microglia in the pathogenesis of AD, they did not address how missense mutations in the protein could modify the risk of developing LOAD with age. Furthermore, a caveat with the interpretation of functional endpoints from AD models crossed with *TREM2* KO mice is that loss of TREM2 function (or its binding partner DAP12) in humans results in Nasu-Hakola disease, a white matter-targeting age- dependent neurodegenerative disease (Koseoglu et al., 2018), suggesting that absence of TREM2 has detrimental effects on the brain in the absence of plaques.

The R47H missense mutation in *TREM2* is strongly and reproducibly linked to LOAD, and since its discovery multiple studies have attempted to model this mutation in mice and rats. Several approaches have produced *TREM2*\*R47H models, including via bacterial artificial chromosome (BAC) and via CRISPR/Cas9 technology. Utilizing a BAC, transgenic mice expressing human common variant (CV) and R47H variants of *TREM2* have been crossed with *TREM2* KO 5xFAD mice, with the resultant phenotype phenocopying *TREM2* KO, therefore suggesting *TREM2*\*R47H variant is a loss-of-function allele of TREM2 (Song et al., 2018). As an extension of this approach, heterozygous human *TREM2*\*R47H and CV cDNA has also been knocked into the mouse *Trem2* locus (Sayed et al., 2021), such that animals express human *TREM2*, but without the full complement of regulatory machinery. Crossing of these mice with the PS19 mouse model of tauopathy found that the R47H variant exacerbates damage and inflammation and does not function as a loss of function variant (Sayed et al., 2021), unlike similar crosses of PS19 mice with the BAC *TREM2*\*R47H model which actually protected against microglia activation and subsequent neurodegeneration (Gratuze et al., 2020), as did crosses with PS19 mice and *TREM2* KO mice (Leyns et al., 2017; Sayed et al., 2018). In addition to these humanized approaches, several *TREM2*\*R47H mouse models have been generated via CRISPR/Cas9 technology and have reported similar findings as *Trem2* KO mice (Cheng-Hathaway et al., 2018; Xiang et al., 2018). However, it has since been reported that introduction of the R47H variant into mouse *TREM2* introduced aberrant splicing, due to synonymous base changes co-introduced with repair templates to arrest Cas9 mediated cleavage. These models display significantly reduced *Trem2* expression (Xiang et al., 2018), effectively making them hypomorphic alleles of *Trem2* that do not accurately reflect the human condition. Understanding how TREM2 and its variants influence the development of LOAD is critical for our understanding of the disease. Therefore, the production of animal models that faithfully reproduce human gene function is crucial to accurately recapitulate the disease in rodents. To that end we, as part of the MODEL-AD consortium, embarked on the development of a *Trem2*\*R47H mouse variant without the shortcomings of artificial cryptic splicing and reduced expression, by using a CRISPR/Cas9 approach utilizing an alternative repair template inspired by Cheng et al., 2018. The resultant *Trem2*^R47H NSS^ mouse has normal *Trem2* expression and normal splicing and is available without restriction to both academic and commercial entities (Jax stock: #034036). To investigate whether correct introduction of the R47H mutation results in a loss of function of TREM2, we utilized the cuprizone model of demyelination to assess the capacity of microglia to clear white matter debris and found that our model shared similar inflammation responses and pattern as wild-type mice, unlike cuprizone treated *Trem2* KO mice. However, despite normal induction of microglial evoked inflammation (including expression of “DAM” genes), we see evidence of increased oligodendrocyte gene expression loss in both *Trem2*^R47H NSS^ and *Trem2* KO mice, consistent with the notion that dysfunctional or absence of TREM2 exacerbates damage in response to a suitable stimulus. Thus, the presence of the R47H variant with normal *Trem2* expression levels does not appear to function as a loss of function allele in response to a cuprizone challenge in terms of a microglial response but does phenocopy the exacerbated damage inferred by the null allele.

To give relevance to AD, we crossed *Trem2*^R47H NSS^ mice with the 5xFAD mouse model of amyloidosis and evaluated pathology and gene expression at 4 and 12 months of age. We identified a consistent sex difference in the initial appearance of plaques, with female *Trem2*\*R47H carriers producing more plaques, but male carriers far less, than their controls. Notably, a similar sex difference has also been observed in *Trem2* KO mice crossed with APP1/PS1 mice ((Delizannis et al., 2021); females have more plaques) and human *TREM2*\*R47H cDNA mice crossed to the PS19 tauopathy model (females have more inflammatory gene expression and spatial memory deficits), as well as transcriptomic analysis of R47H-carrying AD patients ((Sayed et al., 2021); females upregulating immune activation pathways while males upregulate metabolic and adenosine triphosphate pathway). No sex difference was observed by 12 months of age, but plaque density was increased by the presence of the R47H mutation, while both soluble and insoluble Aβ levels are also increased. Consistent with the R47H variant inferring a loss-of-function phenotype, the initial microglia-plaque interaction is impaired, resulting in smaller plaques yet an increase in dystrophic neurites produced by those plaques, in line with prior data from *TREM2* KO mice (Meilandt et al., 2020; Wang et al., 2016; Yuan et al., 2016b). However, these impairments between microglia and plaque interactions are absent by 12 months of age, suggesting a normalization of microglial behavior with time.

Supporting these data, gene expression from microdissected hippocampi mirror the initial impairments between microglia and plaques, with reduced expression of DAM genes such as *Itgax* and *Cd74*. However, by 12 months, when no impairments between microglia and plaques are seen, only selected DAM genes remain reduced – most notably *Itgax*. The presence of the R47H variant induces a selective upregulation of interferon related genes such as as *Ifi47*, *Ifit1- 3*, and *Gbp2*, *6*, and *7*, which are all key players in pathogen response (Li et al., 2017). Furthermore, the WGCNA identified inflammatory module (red) revealed increases in inflammation in the 5xFAD mice at 4 months of age were prevented by the presence of the R47H variant, but equal or exceeding 5xFAD levels by 12 months of age. Thus, the R47H variant appears to confer age and disease specific effects on microglia. Of direct relevance and validating these results, similar findings have been shown in human AD tissue from *TREM2* variant carriers, in which microglial responses to pathology are suppressed in newly formed pathological areas but exacerbated in more advanced pathological brain areas (Prokop et al., 2019).

Given the fact that the R47H *TREM2* variant has been associated with several neurodegenerative diseases we have also focused on how this variant may be more permissive of damage exerted on the brain by the relevant pathology, in this case plaques. As mentioned earlier we see initial increases in dystrophic neurites induced by plaques in the presence of the R47H variant, supporting this notion. We further demonstrate increased plasma NfL, a reliable marker of brain injury that tracks with cortical thinning and cognitive decline in AD populations (Bacioglu et al., 2016; de Wolf et al., 2020; Lee et al., 2022; Quiroz et al., 2020), in the presence of plaques. We localized NfL in the brain to being associated with dystrophic neurites induced by plaques/microglia and found that NfL level in the brain insoluble fraction correlated with levels in the plasma. Collectively, these results highlight how the R47H *Trem2* variant can induce greater damage on clinically relevant endpoints.

We also explored hippocampal LTP and found that the presence of the R47H variant in 5xFAD protected against initial deficits found in the 5xFAD mice, as well as protected against initial loss of pre-synaptic puncta. These lack of LTP deficits and synaptic loss coincided with the impaired microglial response to the plaques, suggesting that TREM2 and microglia have multiple actions that run counter to one another. On one hand, the impaired microglia-plaque interactions promote increased dystrophic neurites and NfL, but also prevent increases in inflammatory gene expression, LTP deficits, and synaptic loss. However, by 12 months of age, LTP deficits were seen in 5xFAD / *Trem2*^R47H NSS^ along with exacerbated presynaptic puncta loss compared to 5xFAD mice. Notably, by 12 months of age *Trem2*^R47H NSS^ mice on a WT background also demonstrated robust impairments in LTP, as well as synaptic puncta loss. Similar results have been reported in a *Trem2*\*R47H rat model, further implicating microglia and dysfunctional *Trem2* in effecting neuronal structure and function (Ren et al., 2020).

A major objective of this study is to provide the scientific community with a reliable and well- characterized mouse *Trem2*\*R47H knock-in model and highlight how this variant has age- and disease- dependent effects on microglia and neuropathology, mirroring reported human data. We demonstrate that *Trem2*\*R47H has profound effects on the synaptic landscape with age, coinciding with robust impairments in LTP. Collectively, these results help to clarify prior data obtained from amyloidosis models crossed with *Trem2*\*R47H mice with unintended hypomorph phenotypes (Cheng-Hathaway et al., 2018; Xiang et al., 2018), and add to our understanding of how microglia and TREM2 contribute to the pathogenesis of AD.

## Acknowledgements

This study was supported by the Model Organism Development and Evaluation for Late-onset Alzheimer’s Disease (MODEL-AD) consortium funded by the National Institute on Aging (U54 AG054349), as well as by R01NS083801 (NINDS), RF1AG056768 (NIA), and RF1AG065329 (NIA) to KNG, and 1F31NS111882-01A1 (NINDS) to MAA. The *Trem2^R47HNSS^* model is available from The Jackson Laboratory (Stock #034036) without restrictions on its use by both academic and commericl users. The content is solely the responsibility of the authors and does not necessarily represent the official view of the National Institutes of Health. We thank Shilpa Sambashivan for providing access to RNA-seq data sets.

---

**Supplemental 1:**
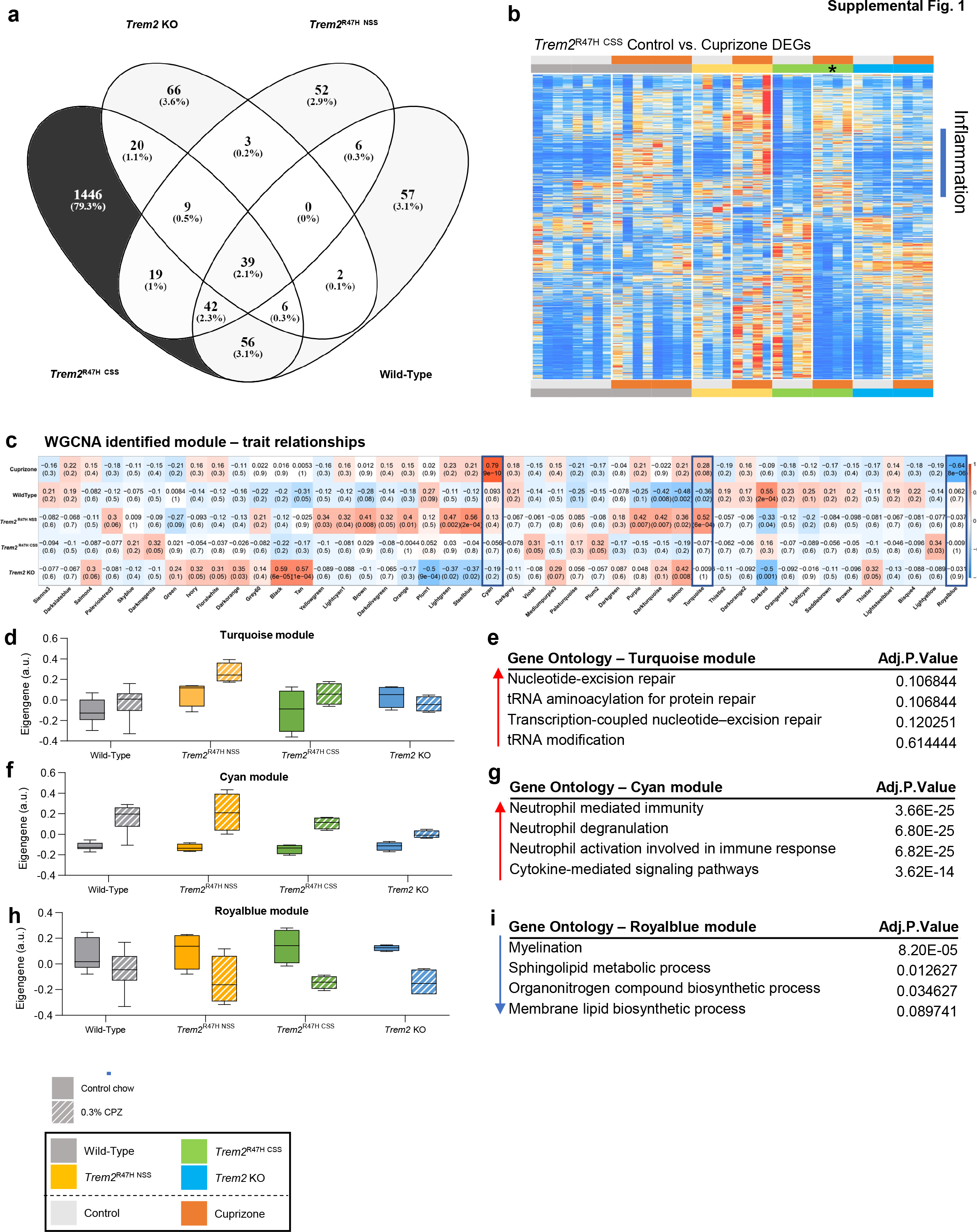
*Trem2*^R47H CSS^ displays a distinct set of differentially expressed genes. **a** Venn diagram displaying number of DEG’s induced only by *Trem2*^R47H CSS^ but not in wild-type, *Trem2*^R47H NSS^ or *Trem2* KO. **b** Heatmap generated from uniquely expressed DEG’s by *Trem2*^R47H CSS^ compared to the other mouse models on control or cuprizone diet. **c** module Trait relationship heatmap by WGCNA on wild-type, *Trem2*^R47H NSS^, *Trem2*^R47H CSS^, and *Trem2* KO associated with cuprizone treatment. Color corresponding to correlation (red color means positive correlation and blue means negative correlation) and the number in parenthesis shows how significance of the correlation. Three modules (turquoise, cyan and royalblue modules) were chosen based on their significant correlation with cuprizone treatment **d, f and h** Barplot for the eigengene of the genes in the turquoise, cyan and royalblue modules respectively . **e, g and i** Gene ontology analysis of the genes in the turquoise, cyan and royalblue modules respectively.

**Supplemental 2:**
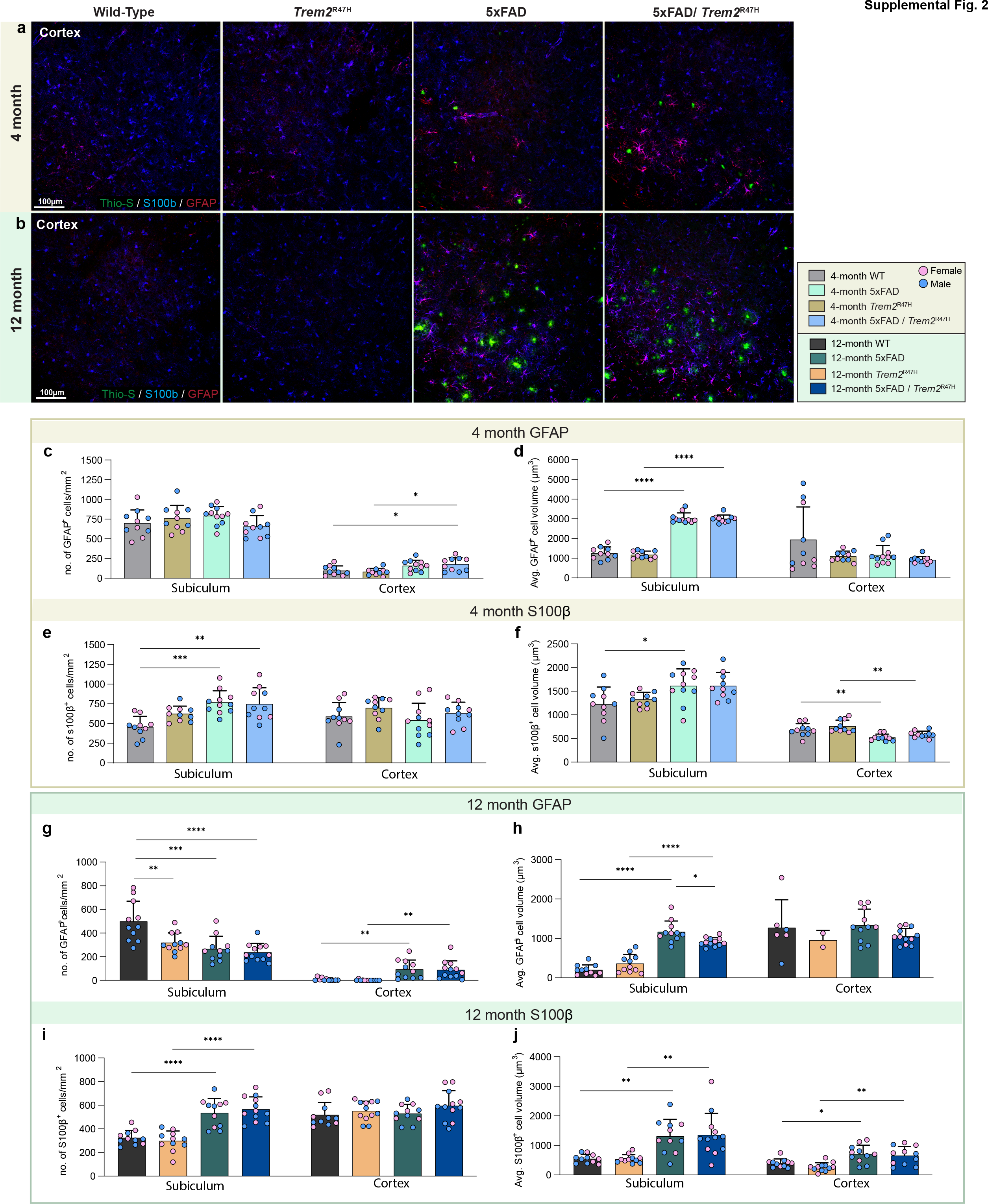
Decrease in astrocyte cell volume in 5xFAD/ *Trem2*^R47H^ at 12-month. **a, b** Representative 20x confocal images of visual cortex stained for dense-core plaques (Thio-S in green), immunolabeled reactive astrocytes (GFAP in red and s100β in blue). **c, d** At 4-month, quantification of GFAP^+^ cell density and average cell volume revealed no difference in cell number but larger cells between 5xFAD and 5xFAD/ *Trem2*^R47H^ and their age-matched controls in subiculum (p<0.0001). Cell number in the visual cortex is higher in 5xFAD and 5xFAD/ *Trem2*^R47H^ compared to their age-matched controls despite no change in cell volume (p<0.05). **e, f** Quantification of s100β ^+^ cell density and average cell volume revealed increase in cell number of 5xFAD compared to WT and larger cells in 5xFAD compared to WT in the subiculum (WT vs 5xFAD, cell no. p<0.001, cell volume p<0.05). In the cortex, there is no difference in s100β ^+^ cell number but smaller cell volume in 5xFAD and 5xFAD/ *Trem2*^R47H^ compared to their controls (p<0.01). **g, h** At 12-month, GFAP^+^ cell number of *Trem2*^R47H^, 5xFAD, and 5xFAD/ *Trem2*^R47H^ mice were reduced compared to WT (WT vs *Trem2*^R47H^, 5xFAD, and 5xFAD/ *Trem2*^R47H^, p<0.01, p<0.001, p<0.0001, respectively) while cell volume is larger in 5xFAD compared to 5xFAD/ *Trem2*^R47H^ (p<0.05). In the cortex, there are more GFAP^+^ cells in 5xFAD and 5xFAD/ *Trem2*^R47H^ compared to controls but no differences in size (p<0.01). **i, j** Quantification of s100β ^+^ cell density and average cell volume revealed increases of both in 5xFAD and 5xFAD/ *Trem2*^R47H^ compared to controls in the subiculum (cell no., p<0.0001; cell volume, p<0.01). In the cortex, there is no difference in s100β ^+^ cell density but an increase in cell volume in 5xFAD and 5xFAD/ *Trem2*^R47H^ (WT vs 5xFAD, p<0.05; *Trem2*^R47H^ vs 5xFAD/ *Trem2*^R47H^; p<0.01). n=10-12. Data are represented as mean ± SEM. Statistical significance is denoted by * p<0.05, ** p<0.01, ***p<0.001, ****p<0.0001.

## Notes

**Conflict of interest:** KNG is a member of the advisory board of Ashvattha Therapeutics

http://swaruplab.bio.uci.edu:3838/5xFAD_Trem2/

### Competing Interest Statement

KNG is a member of the advisory board of Ashvattha Therapeutics

